# Timeline of changes in spike conformational dynamics in emergent SARS-CoV-2 variants reveal progressive stabilization of trimer stalk and enhanced NTD dynamics

**DOI:** 10.1101/2022.08.26.505369

**Authors:** Sean M. Braet, Theresa S. C. Buckley, Varun Venkatakrishnan, Kim-Marie A. Dam, Pamela J. Bjorkman, Ganesh S. Anand

## Abstract

SARS-CoV-2 emergent variants are characterized by increased transmissibility and each show multiple mutations predominantly localized to the spike (S) protein. Here, amide hydrogen/deuterium exchange mass spectrometry has been applied to track correlative changes in S dynamics from multiple SARS-CoV-2 variants. Our results highlight large differences across variants at two loci with impacts on S dynamics and stability. A significant enhancement in stabilization first occurred with the emergence of D614G S followed by smaller, progressive stabilization in Omicron BA.1 S traced through Alpha S and Delta S variants. Stabilization preceded progressive enhancement in dynamics in the N-terminal domain, wherein Omicron BA.1 S showed the largest magnitude increases relative to other preceding variants. Changes in stabilization and dynamics resulting from specific S mutations detail the evolutionary trajectory of S protein in emerging variants. These carry major implications for SARS-CoV-2 viral fitness and offer new insights into variant-specific therapeutic development.

## Introduction

SARS-CoV-2 was first identified in late 2019 and is the causative agent of the ongoing coronavirus pandemic (Chang et al., 2020). Extensive efforts have sparked the development of a number of vaccines and therapeutics to mitigate the effects of infection. However, the emergence of numerous variants of concern have created an additional challenge to treatment and prevention efforts. The periodic emergence of new variants of concern beginning with Alpha, and later including Delta and most recently Omicron BA.1, have contributed to surges in cases worldwide (Wassenaar et al., 2022).

SARS-CoV-2 is a member of the family *Coronavirdae* along with other human pathogens including SARS-CoV and MERS (Corman et al., 2018). The SARS-CoV-2 virion is enveloped and encapsulates a 30 kb +ssRNA genome that encodes envelope (E) protein, membrane (M) protein, spike (S) protein, as well as 16 non-structural proteins and 9 accessory proteins (Ke et al., 2020). S, a critical viral protein for SARS-CoV-2 entry that is targeted by neutralizing antibodies, plays a multifunctional role in the infection process and is therefore a target for vaccine development (Martinez-Flores et al., 2021). S is a glycosylated homotrimer with each monomer consisting of S1 and S2 subunits (Fig. 1A). The S1 domain comprises an N-terminal domain (NTD), a receptor-binding domain (RBD) and two subdomains SD1 and 2, with the RBD mediating the interaction interface with the human ACE2 receptor (Lan et al., 2020). The S2 domain includes the S1/S2 and S2 proteolytic cleavage sites as well as a fusion peptide. During the viral entry process, the S protein is processed by furin protease at the S1/S2 cleavage site either prior to or after S binding to ACE2 receptor. This enables secondary cleavage by a separate protease (commonly transmembrane serine protease 2 (TMPRSS2) or cathepsin) at the S2 site (Peacock et al., 2021; Shang et al., 2020; Vankadari, 2020), which leads to dissociation of S1 and release of the S2 subunit to drive membrane fusion and cellular entry.

**Fig. 1.**
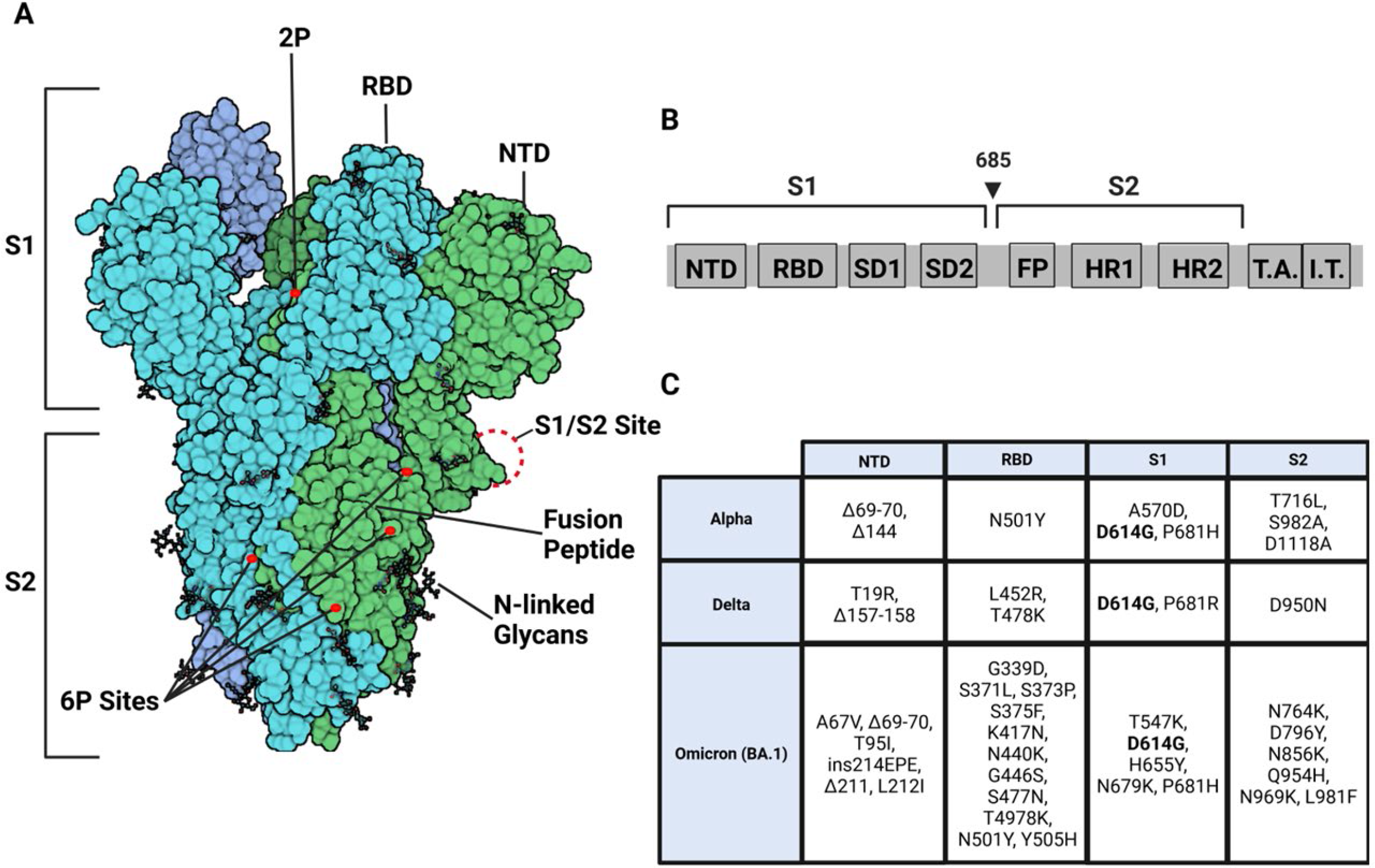
S trimer modifications and variant mutations. **(A)** S trimer (PDB:6VSB) (protomers in dark blue, teal, and green) with RBD, NTD, S1/S2 site, fusion peptide, 2P substitutions (985-986), additional 6P substitutions (817, 892, 899, 942), and glycans. (Image created in Biorender.com) **(B)** Sequence organization for SARS-CoV-2 S protein (NTD = N-terminal domain, RBD – receptor binding domain, SD1 = subdomain 1, SD2 = subdomain 2, FP = fusion peptide, HR1 = heptad repeat 1, HR2 = heptad repeat 2, T.A. = transmembrane domain, I.T. = intracellular domain. The brackets define the recombinant soluble S protein used in this study. Furin cleavage site (685) is indicated by arrow (**C)** Mutations specific to Alpha, Delta, and Omicron BA.1 S variants. D614G is highlighted in bold.

S protein plays three critical roles in facilitating host cell entry: S must bind ACE2, be proteolytically processed, and promote membrane fusion. Domain-specific investigation of S and its variants have provided insights into effects of mutations on functionalities in isolation. However, there is a need to address the composite impact of individual variants on S conformational ensembles in solution (Raghuvamsi et al., 2021). Altered conformations in mutant S proteins from variants would impact interactions of S with ACE2 and downstream functions.

Due to its key roles in viral host recognition and entry, it is unsurprising that S is a hotspot for mutations in emerging variants. A defining feature of emerging variants is each of these became more dominant over the prevailing strains, which has been attributed in part to progressively increased viral fitness (Y. Liu et al., 2022; Plante et al., 2021; Ulrich et al., 2022). Consequently, S D614G showed greater fitness than wild-type S (Plante et al., 2021), the Alpha variant S showed greater fitness than D614G S (Ulrich et al., 2022), and the Delta variant S showed greater fitness than Alpha S (Y. Liu et al., 2022), which likely contributed to surges in human infections. Among the first set of mutations that were detected in S during the early phase of the pandemic, D614G emerged as a dominant variant in 2020 (Chang et al., 2020; Pandey et al., 2021). One of the striking effects observed in a comparison of WT and D614G S proteins revealed a ∼ 50X enhancement in proteolytic processing by furin (Gobeil et al., 2021).

The Alpha variant subsequently emerged in September 2020, and in addition to D614G, the S carried other mutations in NTD (del 69-70, del 144), RBD (N501Y, A570D), the furin-binding site (P681H) and the S2 subunit (T716I, S982A, and D1118H) (Xia et al., 2021) (Fig. 1B). SARS-CoV-2 infections by the Alpha variant were replaced by a more dominant Delta variant, first identified in October 2020. The Delta variant S included mutations in the NTD (T19R, G142D, del 156-157, R158G), RBD (L452R, T478K), furin cleavage site (P681R), and the S2 subunit (D950N) (Tian et al., 2021). A more recent surge in infection has been due to the Omicron variant (BA.1) first identified in November 2021 and was the most highly mutated variant compared to wildtype (Fig. 1B) (L. Liu et al., 2022). Notably, the D614G mutation has been conserved across all major variants of concern (Wassenaar et al., 2022). Additionally, the P681R mutation found in the Delta S and Omicron BA.1 S has been found to increase pathogenicity and proteolytic processing (Y. Liu et al., 2022; Saito et al., 2021), and RBD mutations in variants of concern have been found to increase affinity for the ACE2 receptor (Han et al., 2022; Ozono et al., 2021).

A defining feature of the emergent variants is the apparent progressive increase in transmissibility between humans attributable to corresponding increased viral fitness *in vitro* (Y. Liu et al., 2022; Plante et al., 2021; Ulrich et al., 2022). The basis for increased viral fitness in emerging variants remains poorly understood. Snapshots from single-particle cryo-EM structures of SARS-CoV-2 S trimers have provided structural insights at high resolution (Cai et al., 2020; Duan et al., 2020; Walls et al., 2020; Zhang et al., 2021) but do not completely capture all of the interconverting conformations in solution. Multiple conformations showing RBD in ‘up’ or ‘down’ orientations have been observed by cryo-EM. In the trimer, these translate into closed- all ‘down’ or one, two or all ‘up’ conformations (Barnes, Jette, et al., 2020). These interchanging conformations in solution highlight the ensemble behavior of S. Dynamics of the S ensemble are fundamental to assessing trimer stability and the role of conformational substates in receptor binding, proteolytic processing, and disease propagation. The ensemble properties have been shown to be critical for ACE2 recognition. The RBDs of S protein have been reported to bind the ACE2 receptor only in an ‘up’ conformation (Barnes, Jette, et al., 2020).

Amide hydrogen deuterium exchange mass spectrometry (HDXMS) is a useful method for probing dynamic breathing motions and conformational ensembles in viral systems (Costello et al., 2022; X.-X. Lim et al., 2017; X. X. Lim et al., 2017; Narang et al., 2021; Raghuvamsi et al., 2021). HDXMS uses D_2_O as a probe that labels backbone amides with deuterium dependent upon both solvent accessibility (Peacock et al., 2018) and H-bond propensities (Englander & Kallenbach, 1983). The labeling reaction can be quenched to probe time scales ranging from second to days with shorter timescales impacted primarily by changes in solvent accessibility and longer timescales assessing changes in H-bonding (Peacock et al., 2018). Pepsin proteolysis combined with mass spectrometry provides a readout of deuterium exchange at peptide resolution that can be mapped onto a structure (Hoofnagle et al., 2003). This captures dynamics (> seconds timescale) of the whole ensemble. Further it offers an ability to resolve more than one slow interchanging conformations (if present) by deconvolution of bimodal distributions of deuterium exchanged mass spectral envelopes (Hodge et al., 2020; Hoofnagle et al., 2003; Oganesyan et al., 2018). Decreased deuterium exchange reflects protection from solvent and/or enhanced stability and correspondingly; increased exchange reports increased solvent accessibility and/or disorder.

Comparative HDXMS of recombinant wildtype, D614G, Alpha, Delta, and Omicron BA.1 S variants has allowed us to track changes in intrinsic dynamics across conserved regions of S through the progressively emerging variants of concern. Our results reveal that the timeline of emergence corresponds to an overall stabilizing effect on the S trimer, together with increased dynamics in the NTD and RBD. These loci in the S2 and S1 subunits encompass sites on S showing greatest differences in deuterium exchange in non-glycosylated peptides common across all variants. Peptides showing differential deuterium exchange identified in the trimeric interface referred to as the stalk region in the rest of the study, report inter-protomer interactions while changes in the NTD and RBD report intra-protomer interactions. The D614G point mutation conferred an initial increase in stalk stabilization. This together with increased NTD dynamics in newer variants, contributed to further enhancement of both effects. Timeline analysis reveals that stabilization and enhancement of NTD dynamics effects are independent, with Delta S achieving near maximal stabilization as measured by HDXMS in our experimental timescales. Variants showed varied NTD dynamics with Omicron showing the greatest magnitude increases in NTD dynamics compared to the predecessor variants and wild-type. It is yet to be seen if NTD dynamics can continue to show increases in successive variants of concern. These underscore the coordinated importance of stalk stability together with NTD flexibility upon overall S trimer dynamics, with major consequences for ACE2 recognition, binding, and proteolytic processing.

## Results

### Equilibration at 37 °C (3 h incubation) shifts S ensemble toward prefusion conformation

Recombinant soluble S protein constructs have made them more accessible for structural and biophysical research by obviating the need to culture SARS-CoV-2 viruses, which require extensive safety procedures and related infrastructure. Engineered S ectodomain constructs show increased expression yields aided by ablating furin-like protease cleavage site and enhancing stability through proline substitutions (2P) (Amanat et al., 2021) and 6P (hexapro) (Hsieh et al., 2020). These engineered constructs also eliminated the need for detergent solubilization by excluding the transmembrane C-terminal segments that are embedded in the lipid bilayer in S protein assembled on intact SARS-CoV-2 particles (Barnes, West, et al., 2020).

We carried out our HDXMS analysis with a construct containing either 2P or 6P substitutions and with the four amino acid furin cleavage motif (RRAR) substituted with a single alanine (Amanat et al., 2021). These were expressed in HEK-293T cells (Barnes, West, et al., 2020). S trimers were purified by Ni-NTA and size exclusion chromatography (SEC) as described in methods. The trimer state was independently verified by cryo-EM analysis (Barnes, Jette, et al., 2020; Barnes, West, et al., 2020). S is multiply glycosylated with 22 potential N-linked glycosylation sites (Watanabe et al., 2020). We confirmed our purified trimeric S protein was multiply glycosylated and mapped N-linked glycosylation sites by bottom-up proteomics as described in methods. Of 22 potential N-linked glycosylation sites, we identified 20 sites in wild type S, 21 glycosylation sites in D614G and Delta variant S proteins, and 19 glycosylation sites in Omicron BA.1 S (Table S1). Wild-type and variant S proteins in subsequent sections denote either a 2P (Pallesen et al., 2017) or 6P (hexapro) (Hsieh et al., 2020) as described.

We carried out comparative HDXMS on 2P and 6P engineered wild-type S protein to probe differences in dynamics. 160 non-glycosylated pepsin fragment peptides provided S sequence coverage of 53.1% (Figure 2-figure supplement 1). HDXMS was measured at time points (Dex = 1-10 min). A deuterium exchange difference map and peptide level significance testing (shown in Woods plots (Lau et al., 2021) of the two constructs) revealed no substantial differences (Δ<0.5 Da) (Figure 2-figure supplement 3-4), indicating both 2P and 6P constructs offered a common baseline for assessing differences across S variants.

The soluble S trimer constructs have been observed to show sensitivity to cold temperature treatment (Costello et al., 2022): Negative stain EM (nsEM) analysis following long-term storage at 4 °C revealed heterogeneous S conformations indicative of trimer instability, whereas incubation at 37 °C for 3h was found to recover a well-formed and more homogenous trimeric structure (Edwards et al., 2021). However, longer incubation times were not found to increase the proportion of S in the trimeric state. It is yet to be determined if this cold sensitivity is an inherent property of the intact S protein or is relevant only to the S trimer ectodomain constructs.

To test the effects of temperature optimization on the S trimer ectodomain, we compared HDXMS on wild-type S treated with and without a 3 h incubation at 37 °C after flash freezing and long-term storage at –80 °C. 170 non-glycosylated pepsin fragment peptides provided a primary sequence coverage of 53.6%, and HDXMS was measured at time points (Dex = 1-10 min) (Figure 2-figure supplement 2). A deuterium exchange heat map (% RFU) of wt S shown in Fig. 2A shows higher relative exchange on the outer edges of the trimer compared to the intratrimer core (Fig. 2A). Decreased exchange alone was observed across multiple regions of S and of high magnitude at the trimer interface (Fig. 2B and D). Decreases were most prominent for peptides in trimer stalk region of S (peptides 899-913, 988-998, 1013-1021) and other interprotomer contacts (peptides 553-568 and 32-48), indicative of increased stability following a 3h incubation at 37 °C (Fig. 2C-E, Figure 2-figure supplement 5).

**Fig. 2.**
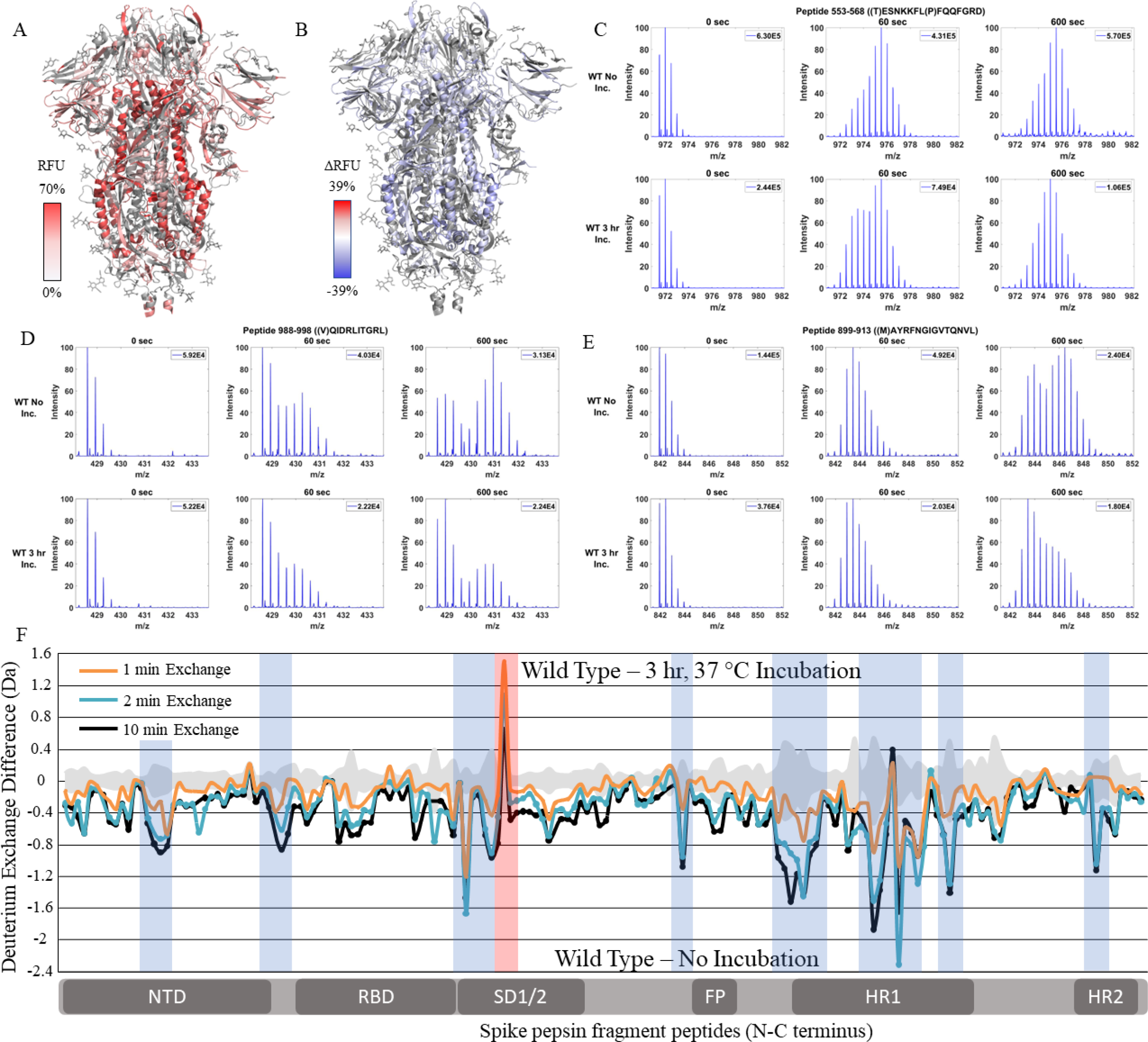
WT S trimer undergoes temperature dependent trimer-monomer transitions. **(A)** Relative fractional uptake (t = 10 min) for unincubated wildtype S protein mapped onto an S trimer structure with three ‘down’ RBDs (PDB ID: 6VXX) (coverage of wildtype spike constructs shown in Figure 2-figure supplement 1-2, differences in deuterium exchange for wildtype 2P and 6P constructs shown in Figure 2-figure supplement 3). White denotes low deuterium exchange and shades of red denote high deuterium exchange. **(B)** Differences in deuterium exchange (ΔRFU) (t = 10 min) for wildtype S protein after a 3 h incubation at 37 °C minus unincubated wild type S protein were mapped onto the S protein structure (PDB ID: 6VXX). Shades of blue correspond to decreased deuterium uptake and shades of red correspond to increased deuterium uptake. **(C-E)** Stacked mass spectra for wildtype S peptides 553-568, 899-913, and 988-998 with undeuterated reference spectra, 1 min and 10 min exchange (left to right). For each peptide, the top row contains spectra for unincubated wild type S and the bottom row contains spectra for wild type S protein that was incubated for 3 h at 37 °C for 3 h. **(F)** Differences in deuterium exchange (deuterons) mapped at peptide resolution from N to C terminus for wild type S incubated for 3 h at 37 °C minus unincubated wildtype S are shown in difference plots for 1, 2, and 10 min exchange. Blue boxes correspond to significantly protected peptides and red boxes correspond to significantly deprotected peptides. Significance was determined by peptide level significance testing (p<0.01, Figure 2-figure supplement 4-5).

Several peptides showed spectral broadening reflective of ensemble behavior in S protein in solution (Table S2). Resolvable bimodal mass spectral distributions were evident for peptides 899-913 (Table 1A) and 988-998 (Table 1B). Bimodal deuterium exchange spectra are attributable to EX1 deuterium exchange kinetics with comparable rates of protein refolding and observed rates of deuterium exchange (k_obs_) (Kaltashov & Eyles, 2002; Weis et al., 2006). We infer that the basis for the bimodal exchange at peptides 899-913 and 988-998 that we observed in our HDXMS experimental timescales are indicative of localized trimer-protomer transitions at the interprotomer interface. Stabilization resulting from 3h 37 ℃ incubation is reversible. Replicate analysis of 37 °C stabilized S trimers with incubation at 4 ℃ prior to deuterium exchange (see methods) showed a time dependent reversal of stabilization as reported previously (Costello et al., 2022), most evident at the same peptides. These results highlight temperature sensitive reversible contacts at the edge of a trimer interface core or trimer stalk region. This region encompasses a long central helical segment (987-1031) and a helix flanking the heptad repeats (900-913) (Walls et al., 2020). **Table 1**.

**Table 1.** Bimodal distributions for mass spectral envelopes of deuterium exchange in peptides 899-913 and 988-998. The left and right centroids are in mass/charge (m/z). The percentages describe the relative abundances of low (left) and high (right) exchanging populations. (A) Peptide 899-913 (B) Peptide 988-998.

|  |  |  |  |  |  |  |
| --- | --- | --- | --- | --- | --- | --- |
| **A** | 899-913 | Time (min) | Left Centroid (m/z) | Right Centroid (m/z) | Left % | Right % |
|  | Unincubated WT | 10 | 844.0 | 846.6 | 33.9 | 66.1 |
|  | Incubated WT | 10 | 844.3 | 847.2 | 59.3 | 40.7 |
|  | D614G | 10 | 843.9 | 845.6 | 90.4 | 9.6 |
| **B** | 988-998 | Time (min) | Left Centroid (m/z) | Right Centroid (m/z) | Left % | Right % |
|  | Unincubated WT | 10 | 429.1 | 430.9 | 37.4 | 62.6 |
|  | Incubated WT | 10 | 429.0 | 430.7 | 62.0 | 38.0 |
|  | D614G | 10 | 643.1 | 644.8 | 90.6 | 9.4 |

### Global conformational changes conferred by the D614G substitution

One of the earliest conserved mutations identified in emergent variants was D614G (Pandey et al., 2021) which demonstrated increased viral fitness along with enhanced furin proteolytic cleavage (Gobeil et al., 2021). To measure the impact of this mutation upon S dynamics, we compared HDXMS of D614G S with wild-type S. Comparative HDXMS between D614G and wild-type S protein (2P constructs) was carried out using our previously established 37 °C temperature incubation (3h) treatment to compare equivalent trimer stabilized states. No peptides spanning the D614G mutation site were identified and therefore all peptides analyzed were common to both wild-type and D614G S. 132 non-glycosylated pepsin fragment peptides were identified covering 47.3% of the D614G sequence (Figure 3-figure supplement 1).

Relative fractional uptake values for the D614G variant mapped onto an S trimer structure (PDB 6VXX) for Dex = 10 min showed a similar relative deuterium exchange profile to that for WT S. The central stalk region showed lower exchange relative to the peripheral surface accessible regions consistent with it forming the stable core of the trimer (Fig. 3A). A difference map (D614G minus wild type) (Dex =10 min) (Fig. 3B) revealed three non-contiguous clusters of peptides distal to D614G site of mutation, showing following differences in exchange: i) Decreased exchange at trimer interface and ii) Increased exchange at NTD and iii) increased exchange in heptad repeat segments.

**Fig. 3.**
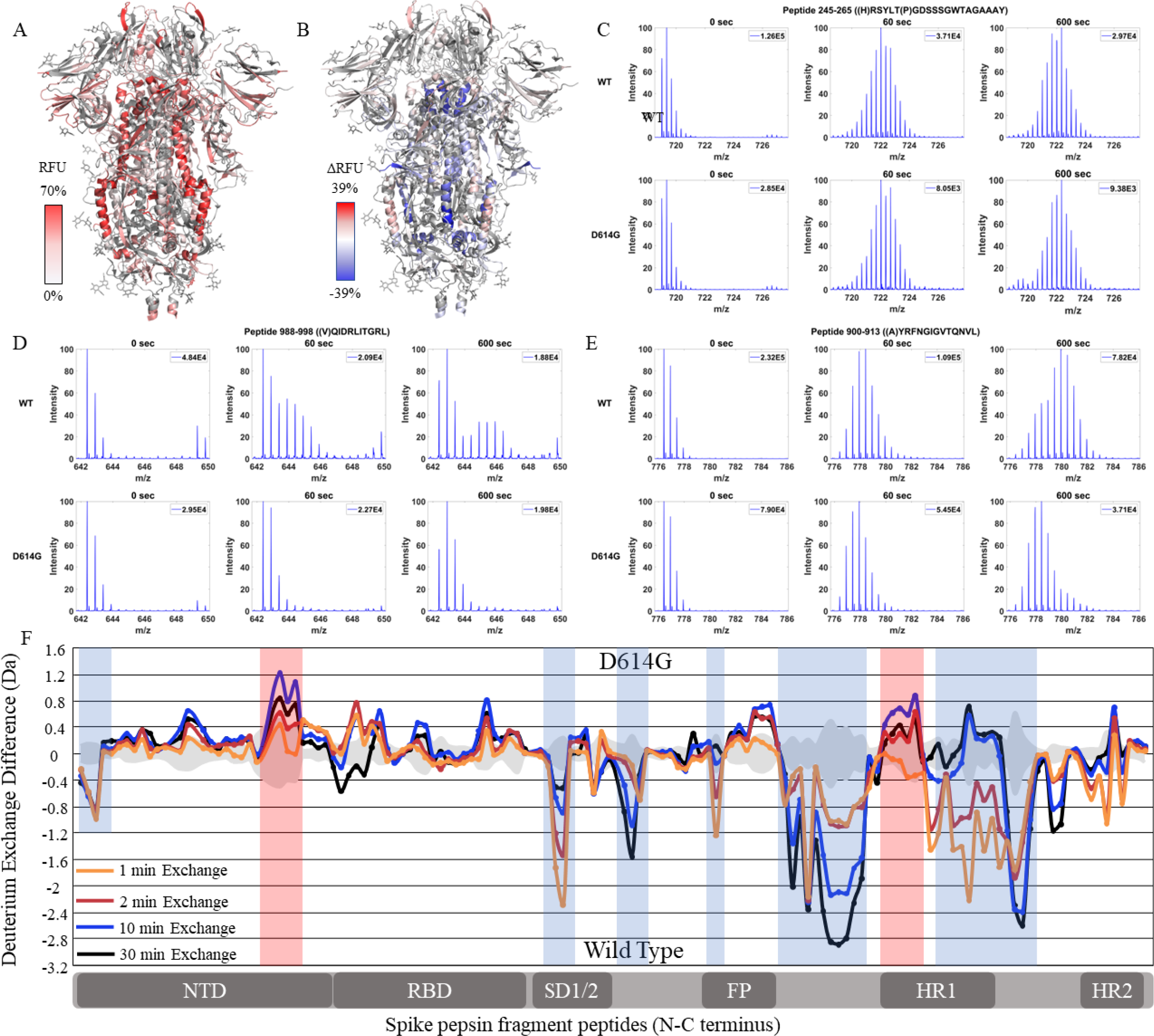
The D614G mutation imparts stabilization on S trimer stalk and leads to increased NTD dynamics. **(A)** Relative fractional uptake (t = 10 min) for D614G S protein mapped onto an S protein structure (PDB ID: 6VXX) (coverage maps shown in Figure 3-figure supplement 1-2, differences in deuterium exchange for D614G S incubated at 37°C minus D614G S without incubation is shown in Figure 3-figure supplement 3). Shades of white correspond to low deuterium exchange and shades of red correspond to high deuterium exchange. **(B)** Differences in deuterium exchange (ΔRFU) (t = 10 min) for D614G S minus wild type S protein were mapped onto an S protein structure (PDB ID: 6VXX). Shades of blue correspond to decreased deuterium uptake and shades of red correspond to increased deuterium uptake. **(C-E)** Stacked mass spectra for wildtype S peptides 245-265, 900-913, and 988-998 with undeuterated reference spectra, 1 min and 10 min exchange (left to right). For each peptide, the top row contains spectra for wildtype S and the bottom row contains spectra for D614G S. **(F)** Differences in deuterium exchange (deuterons) mapped at peptide resolution from N to C terminus for D614G minus wildtype S are shown in difference plots for 1, 2, 10, and 30 min exchange. Blue boxes correspond to significantly protected peptides and red boxes correspond to significantly deprotected peptides. Significance was determined by peptide level significance testing (p<0.01, Figure 3-figure supplement 4). Back exchange for D614G is estimated in Figure 3-figure supplement 5.

### Decreased exchange at the trimer stalk region in D614G

Deuterium exchange difference plots showed a small subset of contiguous peptides from three regions within the trimer stalk region that showed the largest magnitude protection in deuterium exchange (Dex = 1 min) in the D614G variant (Fig. 3C, Figure 3-figure supplement 4). These regions also showed decreased exchange upon a 3 h incubation at 37 °C (Table 2). Representative peptides 899-913, 988-998, and 1013-1021 are reporters for deuterium exchange at the stalk region (Table 2). A similar trend with decreased exchange in this locus with temperature incubation was seen with D614G S as with wild-type S (Figure 3-figure supplement 2-3). Incubating D614G S for 3 h at 37 °C resulted in a smaller degree of stabilization compared to the 3 h 37 °C incubation of WT S (Figure 3-figure supplement 3). These are consistent with stabilization observed previously for D614G (Edwards et al., 2021).

**Table 2.**
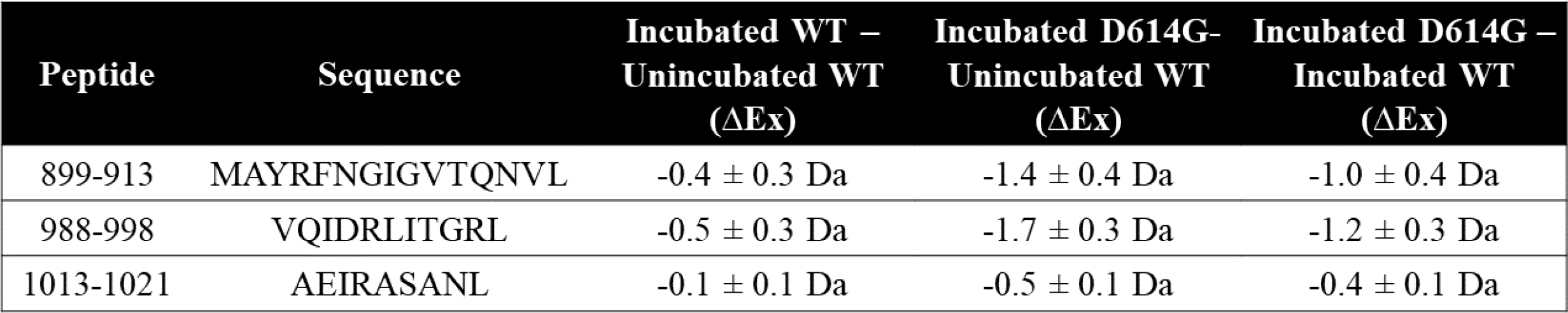
Magnitude of deuterium exchange protection conferred by D614G is greater than that by temperature stabilization. Differences in exchange (Dex =1 min) with temperature incubation for WT compared to D614G mutation for peptides from Spike trimer.

We further observed a spectral broadening of deuterium exchange in stalk peptides in wild-type S and a more resolvable bimodal distribution was observed for peptides 899-913 and 988-998 at 10 and 30 min (table S2). Peptide 899-913 lies near the base of the trimer stalk relative to RBD and exhibits a slow conformational interconversion (time scale ∼ 15-30 min) in which inter-protomer contacts are broken (high exchanging population) and then reassociate (low exchanging population). Consequently, deuterium exchange protection in mutants observed at time points later than D_ex_ = 1 min directly correlates with the rate of reversible localized trimer dissociation (Table 2). The increased strength of interprotomer contacts observed after 1 min deuterium exchange are consistent with slower transitions and a higher proportion of trimer conformations with well-formed interprotomer contacts in S variants. The protection observed after 10 and 30 min exchange is a function of increased inherent stability combined with a shift in ensemble of trimer to favor a more stable, lower exchanging conformation (Table 1). The D614G variant showed lower deuterium exchange overall indicating stronger interprotomer contacts.

### Increased exchange in NTD in D614G S

A striking difference in D614G not found upon 37 °C stabilization of wild-type S was observed in the NTD peptides (peptides spanning regions 177-191 and 243-265) (Fig. 3C, table S3), each of which showed increased exchange relative to wild-type S. This revealed that the single point mutation at D614 to glycine induced long range allosteric effects that are propagated across the trimer and are associated with both stalk stabilization and increased S1 domain dynamics at the NTD. These were the only loci outside the heptad repeats and across the S1 and S2 domains to show significant differences as shown in a Woods plot (p<0.01) (Figure 3-figure supplement 4) in deuterium exchange between wild-type and D614G at non-glycosylated and observed peptides. These effects provided a baseline for tracking conformational changes in S protein in emergent, more transmissible variants. The large conformational changes elicited by the D614G mutation underscore its importance as a highly conserved mutation across emergent variants (Aleem et al., 2022).

**Table 3.** Deuterium Exchange protection at trimer stalk peptides in variants relative to D614G S.

| Peptide | Sequence | Alpha – D614G<br>(ΔEx) | Delta – D614G<br>(ΔEx) | Omicron –<br>D614G (ΔEx) |
| --- | --- | --- | --- | --- |
| 900-913 | AYRFNGIGVTQNVL | -0.1 ± 0.4 Da | -0.3 ± 0.4 Da | -0.3 ± 0.4 Da |
| 990-998 | IDRLITGRL | 0.1 ± 0.3 Da | -0.2 ± 0.3 Da | -0.2 ± 0.3 Da |
| 1013-1021 | AEIRASANL | 0.2 ± 0.2 Da | 0 ± 0.1 Da | 0 ± 0.1 Da |

### Alpha variant S shows increased exchange relative to D614G at both trimer stalk and NTD

We extended our analysis of D614G to variants of concern that each carried this mutation together with multiple other mutations (Fig. 1), comparing each subsequent variant with its epidemiological predecessor to track changes in deuterium exchange across the timeline of emergence. 45.9% coverage was obtained with 127 non-glycosylated pepsin fragment peptides common to D614G, Alpha, and wild-type S for a comparative HDXMS (Dex = 1-30 min) analysis of the Alpha variant S versus D614G S (Figure 4-figure supplement 1). Relative fractional uptake for the Alpha variant was mapped onto a wild type S structure (PDB 6VXX) (Fig. 4A). Differences in deuterium uptake (ΔRFU) for the Alpha variant S minus D614G S are mapped onto PDB 6VXX in Fig. 4B. The Alpha variant S showed lower magnitude changes in deuterium exchange relative to D614G than D614G showed compared to wildtype, particularly in the stalk region. Changes in deuterium exchange were primarily observed at the NTD for common peptides (Figure 4-figure supplement 2).

**Fig. 4.**
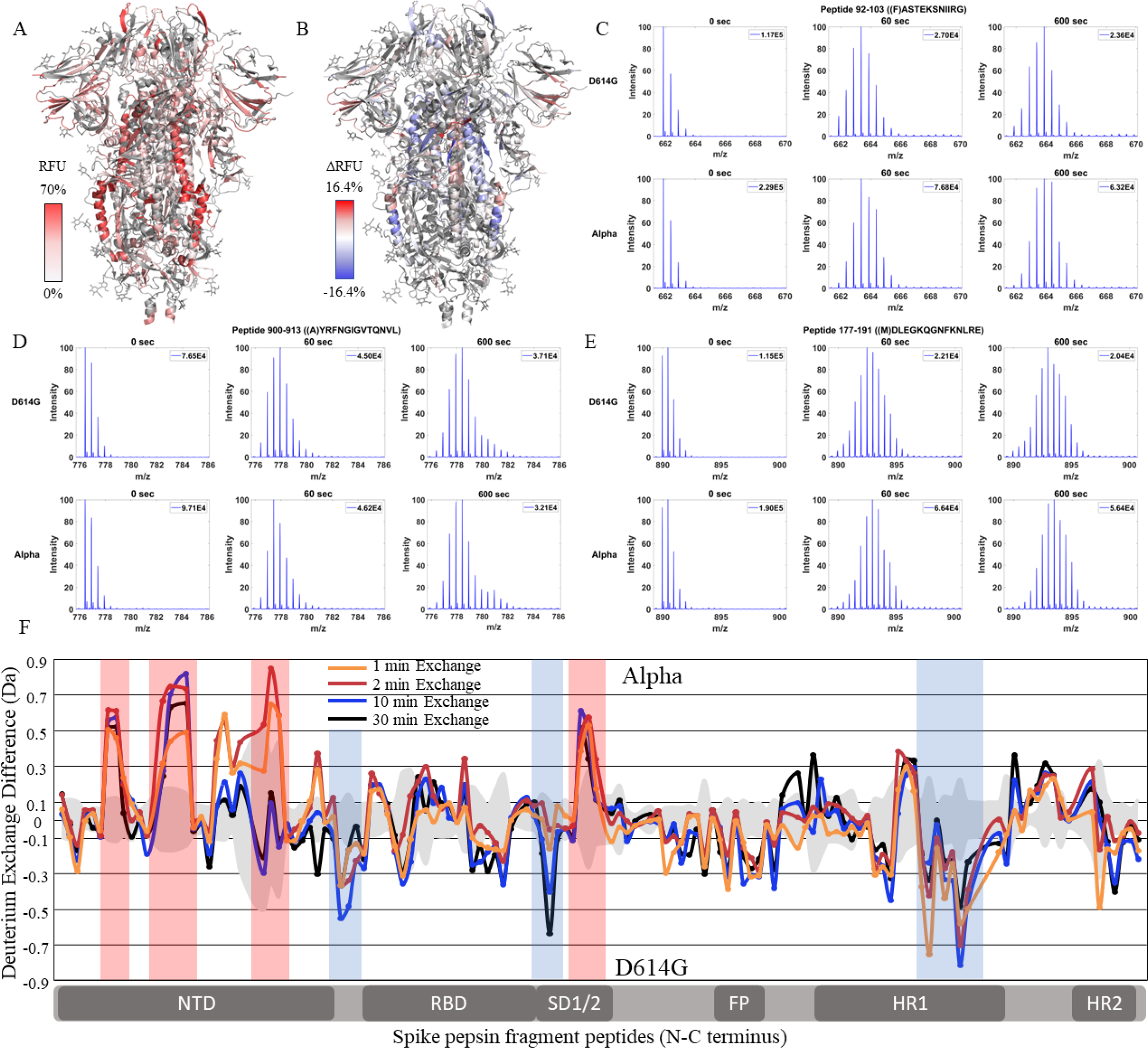
The Alpha S show significantly increased NTD dynamics. **(A)** Relative fractional uptake (t = 10 min) for Alpha S protein mapped onto an S protein structure with three down RBDs (PDB ID: 6VXX) coverage map is shown in Figure 4-figure supplement 1). Shades of white correspond to low deuterium exchange and shades of red correspond to high deuterium exchange. **(B)** Differences in deuterium exchange (ΔRFU) (t = 10 min) for Alpha S protein minus D614G S protein were mapped onto an S protein structure (PDB ID: 6VXX). Shades of blue correspond to decreased deuterium uptake and shades of red correspond to increased deuterium uptake. **(C-E)** Stacked mass spectra for wildtype S peptides 92-103, 177-191, and 900-913 with undeuterated mass spectral envelope as reference, or, 1 min and 10 min exchange (left to right). For each peptide, the top row contains spectra for D614G S and the bottom row contains spectra for Alpha S. **(F)** Differences in deuterium exchange (deuterons) mapped at peptide resolution from N to C terminus for Alpha S minus D614G S are shown in difference plots for 1, 2, 10, and 30 min exchange. Blue boxes correspond to significantly protected peptides and red boxes correspond to significantly deprotected peptides. Significance was determined by peptide level significance testing (p<0.01, Figure 4-figure supplement 2).

In the trimer stalk region for the Alpha S, changes in deuterium exchange in Alpha relative to D614G S proteins were far lower in magnitude than in D614G relative to wildtype in the trimer stalk region. The trimer stalk region in particular showed no differences relative to wild-type S, with peptides 900-913 and 990-998 showing a small magnitude increase in deuterium exchange (0.2-0.3D) (Table 3). This indicated that a bulk of the stabilization of the stalk region in the Alpha S protein was contributed by the conserved D614G mutation.

Outside the stalk, a lower magnitude decrease in exchange was observed in the S2 domain specifically at peptides at the C-terminal end of heptad repeat 1 in the 940-975 region (Δ Ex= 0.4- 0.8 D) as well as peptide 553-568 (Δ Ex = 0.6 D), which mediates inter-monomer contacts. Interestingly, the largest changes were observed in the NTD. The Alpha S showed increased exchange at peptides spanning 92-103, 177-191, and 201-264 (table S3).

### Delta variant S shows decreased exchange at both the trimer stalk and NTD

Comparative HDXMS of the Delta variant S to the Alpha S generated 47.0% coverage was obtained with 123 non-glycosylated pepsin fragmentation peptides common to D614G and the Delta and Alpha S variants (Figure 5-figure supplement 1). RFU for Dex = 1 min in the Delta variant and differences in exchange for Delta S minus Alpha S (Δ RFU) were mapped onto an S structure (PDB: 6VXX) (Fig. 5A, B). Delta S showed mostly decreased exchange relative to the Alpha S with decreases primarily in the trimer stalk, other S2 domain peptides, and the NTD (Fig. 5C-E). Delta S showed decreased exchange in trimer stalk peptides relative to Alpha S (Table 3). In the S2 domain, additional decreases in exchange were observed at regions 820-830 (ΔEx = 0.7D) corresponding to the fusion peptide and 900-940 (ΔEx = 1.2D) corresponding to the N-terminal end of heptad repeat 1. Based on Woods plots analyses (p<0.01), insignificant differences in deuterium exchange were observed for other S2 peptides relative to Alpha (Figure 5-figure supplement 2).

**Fig. 5.**
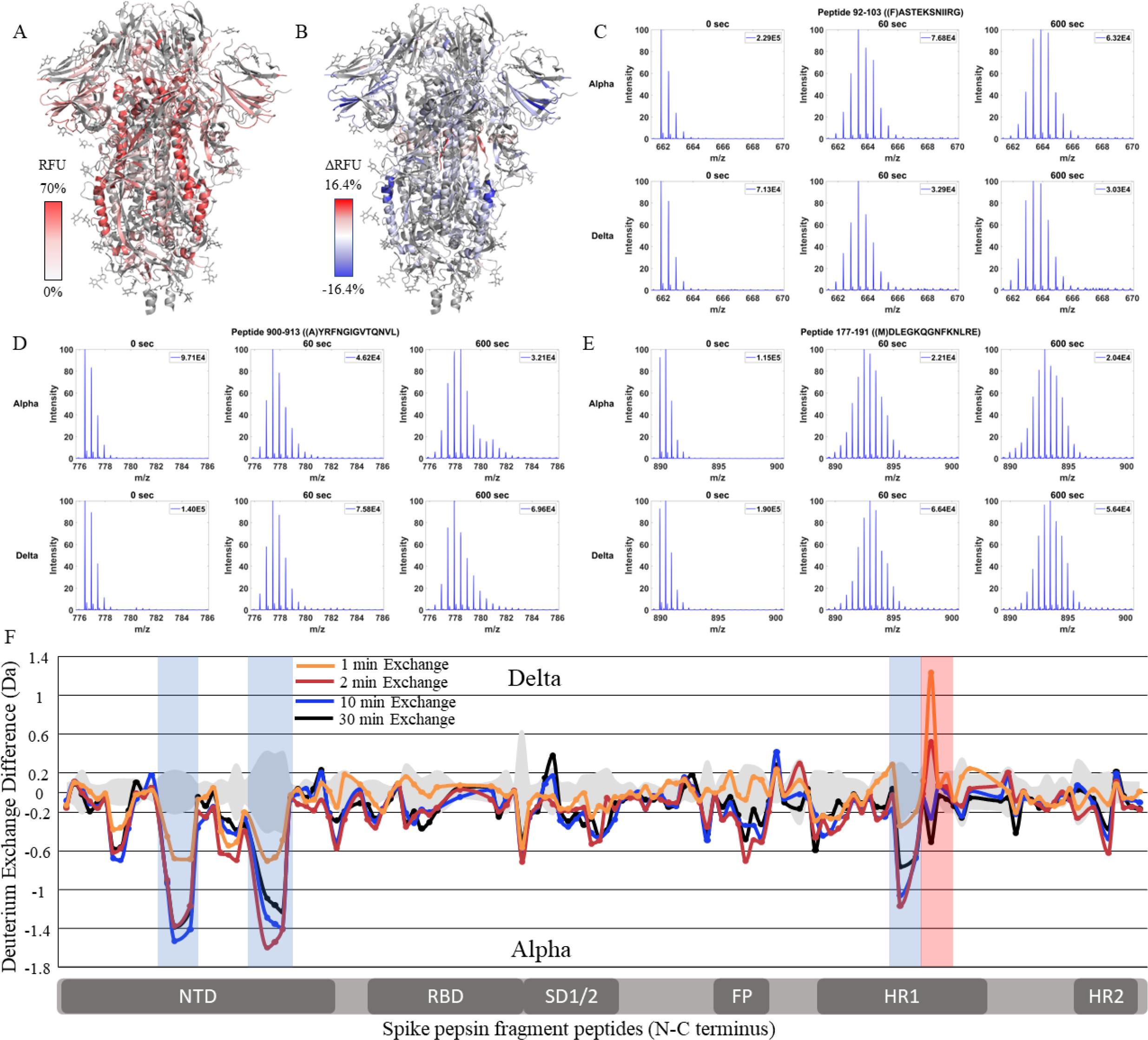
The Delta variant show increased stability with decreased NTD dynamics. **(A)** Relative fractional uptake (t = 10 min) for Delta S protein mapped onto an S protein structure with three down RBDs (PDB ID: 6VXX) coverage maps shown in Figure 5-figure supplement 1). Shades of white correspond to low deuterium exchange and shades of red correspond to high deuterium exchange. **(B)** Differences in deuterium exchange (ΔRFU) (t = 10 min) for Delta S minus Alpha S were mapped onto an S protein structure (PDB ID: 6VXX). Shades of blue correspond to decreased deuterium uptake and shades of red correspond to increased deuterium uptake. **(C-E)** Stacked mass spectra for wildtype S peptides 92-103, 177-191, and 900-913 with undeuterated reference spectra, 1 min and 10 min exchange (left to right). For each peptide, the top row contains spectra for Alpha S and the bottom row contains spectra for Delta S. **(F)** Differences in deuterium exchange (deuterons) mapped at peptide resolution from N to C terminus for Delta S minus Alpha S are shown in difference plots for 1-, 2-, 10-, and 30-min exchange. Blue boxes correspond to significantly protected peptides and red boxes correspond to significantly deprotected peptides. Significance was determined by peptide level significance testing (p<0.01, Figure 5-figure supplement 2).

Delta S also showed decreased exchange for NTD peptides relative to Alpha S. Decreased exchange was most prominent at peptides spanning 92-103, 177-191, and 200-265 (Table S3). Additional decreases in exchange were observed at 306-317 (ΔEx = 0.6D) while increased exchange was observed at the RBD peptide 453-467 (ΔEx = 0.6D). Incremental decreases in exchange at the trimer stalk were in addition to a bulk stabilization imparted by the D614G mutation while decreased exchange in the NTD represented a reversal of the increased NTD exchange observed in Alpha S.

### Omicron BA.1 S variant retains low exchange at the trimer stalk while showing increased deuterium exchange at NTD peptides

Finally, we compared Omicron BA.1 S to Delta S. 36.4% coverage was achieved with 98 non-glycosylated pepsin fragment peptides common to D614G, Delta and Omicron BA.1 S proteins (Figure 6-figure supplement 1). RFU for Dex = 1 min in Omicron BA.1 S and differences in exchange for Omicron BA.1 S minus Delta S (Δ RFU) were mapped onto an S structure (PDB: 6VXX) (Fig. 6A, B). It should be noted that the Omicron BA.1 variant used for analysis was the 6P construct, necessitated by poor expression and heterogeneity of the 2P construct of Omicron BA.1 S. Based on our analysis of wildtype S, the 2P and 6P constructs showed no differences in HDXMS. The Omicron BA.1 S showed slightly decreased exchange at the trimer stalk with increased exchange at other S2 domain peptides, and increased exchange in the NTD (Fig. 6C-E).

**Fig. 6.**
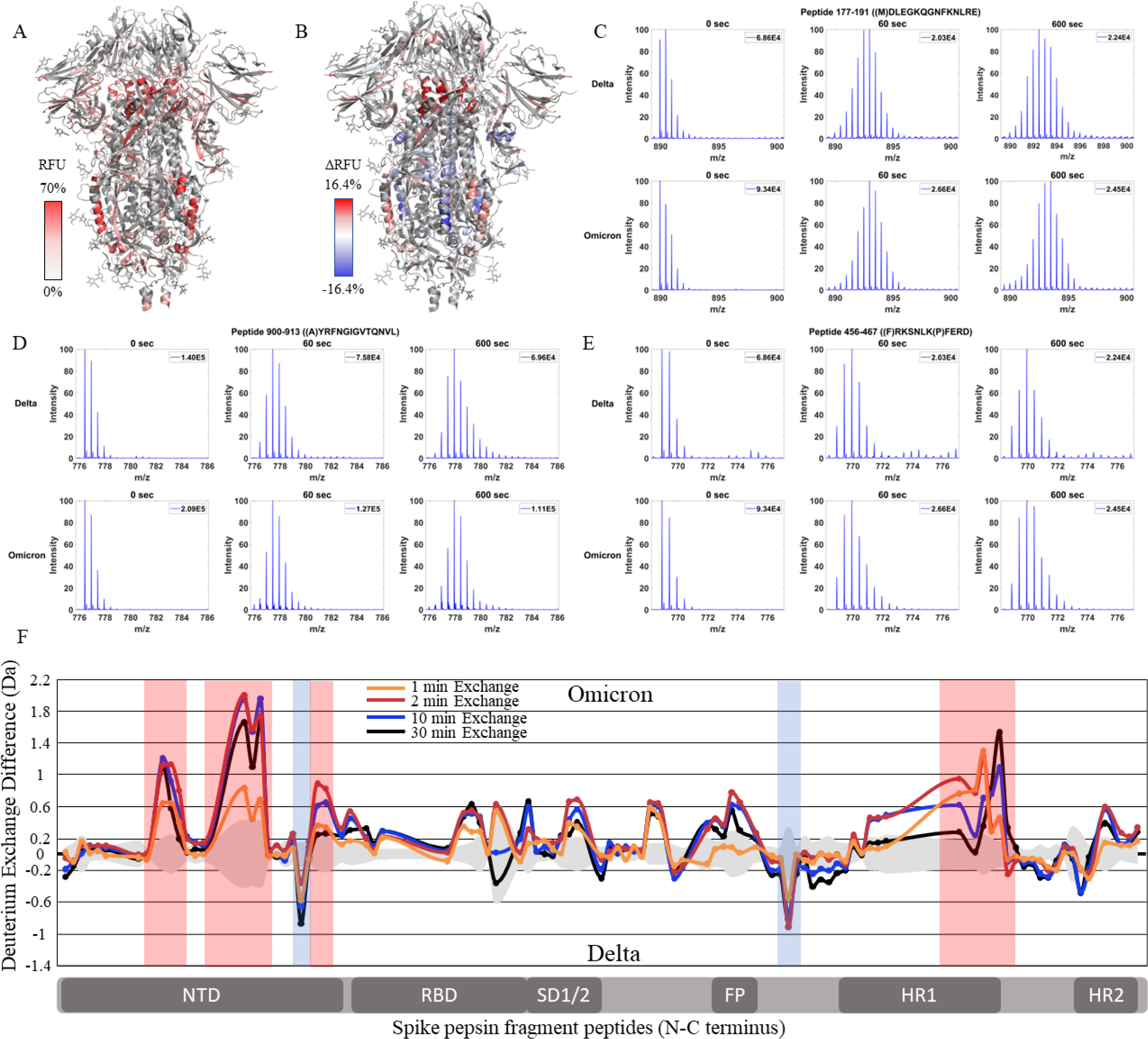
The Omicron BA.1 S shows high magnitude trimer stability and NTD dynamics. **(A)** Relative fractional uptake (t = 10 min) for Omicron BA.1 S mapped onto an S protein structure with three down RBDs (PDB ID: 6VXX) coverage maps shown in Figure 6-figure supplement 1). Shades of white correspond to low deuterium exchange and shades of red correspond to high deuterium exchange. **(B)** Differences in deuterium exchange (ΔRFU) (t = 10 min) for Omicron BA.1 S minus Delta S were mapped onto an S protein structure (PDB ID: 6VXX). Shades of blue correspond to decreased deuterium uptake and shades of red correspond to increased deuterium uptake. **(C-E)** Stacked mass spectra for wildtype S peptides 92-103, 177-191, and 900-913 with undeuterated reference spectra, 1 min and 10 min exchange (left to right). For each peptide, the top row contains spectra for Delta S and the bottom row contains spectra for Omicron BA.1 S. **(F)** Differences in deuterium exchange (deuterons) mapped at peptide resolution from N to C terminus for Omicron BA.1 S minus Delta S are shown in difference plots for 1-, 2-, 10-, and 30-minute exchange. Blue boxes correspond to significantly protected peptides and red boxes correspond to significantly deprotected peptides. Significance was determined by peptide level significance testing (p<0.01, Figure 6-figure supplement 2).

A small additional decrease in exchange was observed for peptides spanning the trimer stalk (Table 3). Notably, the Omicron BA.1 S showed a decrease in the higher exchanging population for stalk peptides suggesting both an impact on inherent trimer stability and ensemble behavior. In S2 domain peptides spanning 920-988 (ΔEx = 0.5-1.5D), 818-830 (ΔEx = 0.4-0.8D), and 750-756 (ΔEx = 0.6D), increased exchange was observed (Figure 6-figure supplement 2).

In the NTD, significantly increased exchange was observed in Omicron BA.1 S relative to Delta S. Increases were seen at peptides spanning 177-191, 243-265, and 306-317 (table S3). Additional increases in exchange were observed for RBD peptide 456-467 (ΔEx = 0.5D) and peptides 553-568 (ΔEx = 0.5D) and 627-643 (ΔEx = 0.5D). Omicron BA.1 S was distinct from the Alpha and Delta S proteins in that it adhered to the continued trend of decreased exchange at the trimer stalk and increased exchange in the NTD that were first observed in D614G S.

## Discussion

### Changes in S conformation: Implications for viral fitness

We report two uncorrelated effects of mutations upon the conformational dynamics of variant S proteins. One of the striking observations from our comparative HDXMS analysis is the progressive stabilization of the S trimer with successive emergent S variants from D614G through Alpha and Delta and subsequently to Omicron BA.1 at the S trimer stalk region. Ensemble behavior is clearly evident from HDXMS of 3 representative peptides from wild-type S. The ensemble shifted to favor a more stable conformation upon incubation at 37 °C. Interestingly, the sequences of the 3 peptides examined in the stalk region adopt an amphipathic helical fold (Figure 7-figure supplement 1). Cold sensitivity followed by spike stabilization at higher temperatures (37°C) can be attributed to the hydrophobic interactions at the stalk region that contribute to stability at the trimer interface (Costello et al., 2022; Edwards et al., 2021; Privalov, 1990).

A first mutation to confer a large stabilization effect was D614G, a lynchpin S mutation, which appeared early during the pandemic and is conserved across nearly all recent emergent SARS-CoV-2 variants of concern (Pandey et al., 2021). Stabilization (associated with decreased exchange) at this stalk locus region showed a leveling off with Delta S (Figure 7-figure supplement 2). Comparative analysis of peptides encompassing mutations in variant S with D614G S further validated D614G being the most prominent contributor to enhanced trimer stabilization (Table S4). However, enhancement in NTD/RBD dynamics appeared more pronouncedly in Omicron BA.1 S protein. Importantly, incubation at 37 ℃ generated a stabilization in the stalk region but did not elicit any changes at the NTD/RBD, indicating that stalk stabilization and enhancement of NTD/RBD dynamics are uncorrelated conformational effects.

Early studies on spike trimers from HIV-1, MERS, and SARS-CoV (Derking & Sanders, 2021; Kirchdoerfer et al., 2018; Pallesen et al., 2017)highlighted analogous stem loci to comprise a conformational dynamic switch to toggle from prefusion to postfusion conformations. To overcome the conformational heterogeneity in SARS-CoV and MERS spike trimers, 2 consecutive proline substitutions were shown to confer improved expression of prefusion trimers (Kirchdoerfer et al., 2018; Pallesen et al., 2017). These mutations were extended to SARS-CoV-2 S protein. The SARS-CoV-2 2P constructs were also found to be more immunogenic, making it a preferred construct for vaccine development (Hsieh et al., 2021; Lien, Kuo, et al., 2021; Lien, Lin, et al., 2021). Mutations introduced on a 2P background were screened for improved expression and 4 additional proline substitutions were identified. These formed the basis for the hexapro (6P) (Hsieh et al., 2020) substitution construct also widely used for structural and biophysical characterization of S. Interestingly, one of the loci where we observed the biggest stabilization in D614G and successive emergent variants is at this locus. A peptide directly C-terminal to the 2P substitutions, 988-998, showed ensemble behavior across S proteins from all variants examined in this study. Omicron BA.1 S showed the highest stabilization at this locus. These independent effects of structure-guided mutations and effects from emergent variants confirm an evolutionary advantage that stabilization of the S trimer stalk region conferred SARS-CoV-2.

### Increasing transmissibility (viral fitness) correlates with more stable trimer and dynamic NTD

Increased transmissibility is a defining feature of S across SARS-CoV-2 variants, with the Omicron BA.1 showing the fastest rate of infection spread (Araf et al., 2022; Y. Liu et al., 2022; Plante et al., 2021; Ulrich et al., 2022). It should be noted that increased transmissibility as assessed from the above referenced *in vitro* studies examining infection of cell lines does not linearly extrapolate to transmissibility in human populations. Comparative HDXMS revealed large scale changes in S conformation across variants. We surmise that these conformational changes would majorly impact multiple functions of the S protein including ACE2 recognition and binding, proteolytic processing by furin-like proteases and TMPRSS2, and efficiency of membrane fusion. The conformational changes identified from changes in deuterium exchange encompassed increased compactness of a central stalk region and variable changes across NTD and RBD regions. Increased transmissibility could be attributable at least in part to a combination of the above conformational changes observed across S variants.

Comparative HDXMS of S variants relative to wild-type S at conserved amino acids revealed that the largest magnitude decreases in exchange were entirely localized to the S2 subunit (Fig. 3). One of the striking features are three key pepsin fragment peptides that are structurally non-contiguous and showed the largest decreases in exchange as well as a progressive shift in ensemble behavior that correlate with timeline of emergence. These 3 peptides spanning the stalk region represent critical trimer interface loci for maintaining a canonical prefusion conformational ensemble (Fig. 7A). Each of these 3 peptides showed differences in magnitude of protection across variants relative to wild-type S in the timescales of HDXMS (t=1- 30 min) (Fig. 7B).

**Fig. 7.**
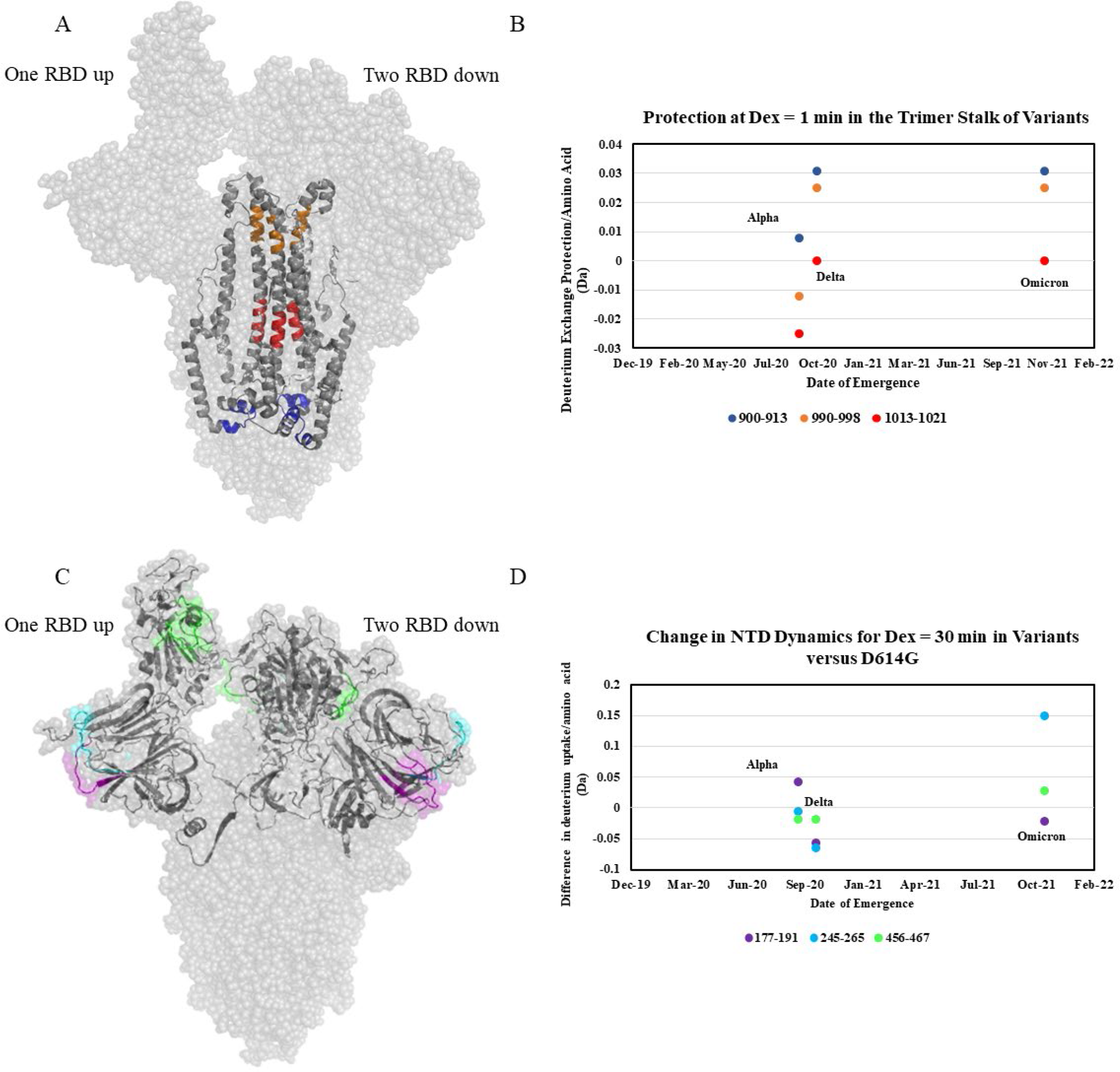
Increased trimer stability and NTD dynamics correlate with the timeline of emergence. **(A)** Trimer stalk peptides 900-913 (blue), 990-998 (orange) and 1013-1021 (red) mapped onto a wildtype S protein structure (PDB ID: 7TGX) (helical wheel analysis of stalk peptides shown in Figure 7-figure supplement 1). **(B)** Protection in trimer stalk peptides in S variants compared to D614G S plotted as protection per amino acid versus date of emergence at Dex = 1 min. **(C)** NTD and RBD peptides showing increased dynamics in the timeline of variant emergence mapped onto a 1 RBD ‘up’ wildtype S structure (PDB ID: 7TGX). Peptides 177-191, 245-265, and 456-467 are shown in purple, cyan, and green, respectively. **(D)** Changes in deuterium uptake for NTD and RBD peptides in variant S proteins compared to D614G S at Dex = 30 min are plotted as change in deuterium uptake versus date of emergence (additional plots in Figure 7-figure supplement 2).

HDXMS clearly demonstrates that not all mutations elicit equivalent changes in conformation/stability, some mutations have disproportionately larger impacts on overall S and trimer conformational interconversion and stability. However, non-linear improvements in trimer stability beyond that conferred by the D614G mutation likely contribute to improved efficacy in S trimer packaging on the SARS-CoV-2 virion. Due to the multiplicative effects of trimer stability on S functions, we postulate that the enhanced stability beyond that conferred by the D614G mutation could account for some of the increased viral fitness observed in Delta and Omicron BA.1 variants (Fig. 7B).

Additionally, we mapped increases in dynamics at the NTD and RBD in the S1 subunit, with Omicron BA.1 S showing the highest overall NTD and RBD dynamics (Fig. 7 C and D).

These changes are likely associated with the role of NTD and RBD dynamics in promoting improved ACE2 recognition and correspondingly increased viral entry across successive variants (Qing et al., 2021). Increased NTD dynamics might facilitate RBD ‘up’ transitions. This, coupled with mutations at the S-ACE2 interface would then facilitate increased efficacy of SARS-CoV-2 host entry. It should be noted that HDXMS reports an average deuterium exchange for an ensemble of conformations when the rates of conformational interconversion are faster relative to rates of exchange at relevant experimental conditions (Hoofnagle et al., 2003). HDXMS results mapped onto a single endstate cryo-EM structure are useful in identifying dynamic loci on the S protein. However, this leads to a distorted view of uniform changes in exchange average across conformational changes throughout the S trimer, since the endstate structure does not represent the entire conformational ensemble. HDXMS of S protein under the experimental conditions and timescales described here reports an average of up and down transitions that cannot be resolved.

In summary, our results localize the impacts of mutations on conformational dynamics to two specific loci: the trimer stalk region and NTD. These offer a timeline of variant emergence effects on S trimer conformational dynamics. Every successive variant S protein displays a progressive reduction in dynamics at a central stalk region in the S2 subunit and progressive increases in dynamics in NTD and RBD. The increased stability conferred by the D614G S substitution formed the basis for further optimization of S conformation through changes in NTD and heptad repeat dynamics that could have advanced overall viral fitness and transmissibility. Alpha S showed no differences at the stalk region but showed increases in NTD and RBD dynamics. Delta S showed more stabilization at the stalk region and decreased NTD dynamics. Omicron BA.1 S showed even greater stabilization of the stalk region together with increased NTD and RBD dynamics. Overall, these results suggest that while near-maximal trimer stalk stability has been achieved, emerging variants continue to show progressive increases in NTD and RBD dynamics. It remains to be seen how progressive changes in stalk stabilization and enhanced NTD dynamics impact neutralization of emergent variants by antibodies generated against wild-type S. Changes in stabilization and dynamics resulting from specific spike mutations detail the evolutionary trajectory of spike protein in emerging variants. This provides a basis for progressively enhanced viral fitness and carries major implications for spike evolution and neutralization.

## Materials and Methods

### Expression and purification of SARS-CoV-2 S proteins

Expression and purification of SARS-CoV-2 S ectodomains were performed as previously described (Barnes, Jette, et al., 2020; Barnes, West, et al., 2020). SARS-CoV-2 S constructs were composed of residues 16–1206 of the early SARS-CoV-2 isolate (GenBank MN985325.1), Alpha variant (GISAID EPI_ISL_601443), Delta variant (GenBank QWK65230.1), or Omicron variant (BA.1) (GISAID EPI_ISL_9845731) with the following stabilizing mutations: 2P (Pallesen et al., 2017) or 6P (Hsieh et al., 2020), the furin cleavage site mutated to Ala, a C-terminal TEV protease site (GSG-RENLYFQG), foldon trimerization motif (GGGSG-YIPEAPRDGQAYVRKDGEWVLLSTFL), 8x-His tag (G-HHHHHHHH), and AviTag (GLNDIFEAQKIEWHE). All S constructs were expressed using the Expi293T transient transfections system (GIBCO). S trimers from clarified transfected cell supernatants were purified over HisTrap High Performance columns (Cytiva), followed by size-exclusion chromatography (SEC) using a Superose 6 increase 10 300 column (Cytiva). Fractions corresponding to S trimers were collected and concentrated in 10% glycerol TBS (20 mM Tris, 150 mM NaCl, pH 8.0) then flash frozen in liquid nitrogen and stored at -80 °C. Downstream purification was completed within 12 h of the transfected cell harvest to maximize the quality of trimeric, well-folded S trimers.

### Bottom-up proteomics and glycan profiling

Recombinant S protein variants were digested with trypsin overnight. Samples were separated by RP-HPLC using a Thermo Scientific EASY-nLC™ 1200 UPLC system connected to a Thermo Scientific™ PepMap C18 column, 15 cm × 75 µm over a 90 min 5-25%, 15 min from 40-95 % gradient (A: water, 0.1% formic acid; B: 80 % acetonitrile, 0.1% formic acid) at 300 nL/min flow rate. The samples were analyzed on the Thermo Scientific™ Orbitrap Eclipse™ Tribrid™ mass spectrometer using DDA FT HCD MS2 method. FT MS1 was acquired at resolution settings of 120K at m/z 200 and FTMS2 at resolution of 30K at m/z 200.

The Thermo Scientific™ Proteome Discoverer™ 2.5 software with the Byonic™ search node (Protein Metrics) were used for glycopeptide data analysis and glycoform quantification. Data were searched against a database containing the Uniprot/SwissProt entries of the model proteins with/out common contaminants and 57 human plasma glycans with a 1% FDR criteria for protein spectral matches. The peptide spectra were also manually validated to confirm identification of glycosylation sites.

### Deuterium exchange

Labeling buffer was prepared by diluting 20X PBS in H_2_O in D_2_O (99.9%). 3 μL of sample were added to 57 μL of labeling buffer for a final labeling concentration of 90.16%. Deuterium labelling was carried out for 1, 2, 10, 30, and 100 min at 20°C using a PAL-RTC (Leap) autosampler. During automated HDXMS experiments protein samples were stored at 0 degrees and stability was assessed by staggering technical replicates. After labelling, equivalent volumes of labeling reaction and prechilled quench solution (1.5 M GndHCl, 0.25 M TCEP) was added to bring the reaction to pH 2.5.

### Mass spectrometry and peptide identification

Approximately 8-10 pmol of sample were loaded onto a BEH pepsin column (2.1 x 30 mm) (Waters, Milford, MA) in 0.1% formic acid at 75 μL/min. Proteolyzed peptides were trapped in a C18 trap column (ACQUITY BEH C18 VanGuard Pre-column, 1.7 µM, Waters, Milford, MA). Peptides were eluted in an acetonitrile gradient (8-40%) in 0.1% formic acid on a reverse phase C18 column (AQUITY UPLC BEH C18 Column, Waters, Milford, MA) at 40 μL/min. All fluidics were controlled by nanoACQUITY Binary Solvent Manager (Waters, Milford, MA). Electrospray ionization mode was utilized and ionized peptides were sprayed onto a SYNAPT XS mass spectrometer (Waters, Milford, MA) acquired in HDMS^E^ Mode. Ion mobility settings of 600 m/s wave velocity and 197 m/s transfer wave velocity were used with collision energies of 4 and 2 V were used for trap and transfer, respectively. High collision energy was ramped from 20 to 45 V while a 25 V cone voltage was used to obtain mass spectra ranging from 50-2000 Da (10 min) in positive ion mode. A flow rate of 5 µL/min was used to inject 100 fmol/μL of [Glu^1^]- fibrinopeptide B ([Glu^1^]-Fib) as lockspray reference mass.

Peptides of wild-type and SARS CoV-2 variant S proteins were identified through independent searches of mass spectra from the undeuterated samples in two steps. First, peptides common to wild-type and variant S proteins were identified from a database containing the amino acid sequence of wild-type and D614G S using PROTEIN LYNX GLOBAL SERVER version 3.0 (Waters, Milford, MA) software in HDMS^E^ mode for non-specific protease cleavage. Search parameters in PLGS were set to ‘no fixed or variable modifier reagents’ and variable N-linked glycosylation.

Deuterium exchange was quantitated using DynamX v3.0 (Waters, Milford, MA) with cutoff filters of: minimum intensity = 2000, minimum peptide length = 4, maximum peptide length = 25, minimum products per amino acid = 0.2, and precursor ion error tolerance <10 ppm. Three undeuterated replicates were collected for wild-type and variant S proteins, and the final peptide list includes only peptides that fulfilled the above-described criteria and were identified independently in at least 2 of the 3 undeuterated samples.

In the second step, the workflow was repeated to identify peptides unique to S protein variants. Pepsin fragment peptides from each variant S protein were identified from a database containing the amino acid sequence of the corresponding variant S. Deuterium exchange in these peptides were analyzed using DynamX 3.0 with identical parameters described above.

### Hydrogen-Deuterium Exchange Analysis

The average number of deuterons exchanged in each peptide was calculated by subtracting the centroid mass of the undeuterated reference spectra from each deuterated spectra. Peptides were independently analyzed for quality across technical replicates. Relative deuterium exchange and difference plots were generated by DynamX v3.0. Relative deuterium exchange plots are reported as RFU which is the ratio of exchanged deuterons to possible exchange deuterons. Back exchange estimates were determined by RFU values from a 24 h labeling experiment and are shown in Figure 3-figure supplement 5. Deuteros (Lau et al., 2021) was used to generate Woods plots using a peptide level significance test (p<0.01). The mass spectrometry proteomics data will be deposited to the ProteomeXchange Consortium via the PRIDE partner repository.

## Funding

Startup funding from the Pennsylvania State University (PSU) to GSA.

We thank Rosa Viner (Thermo Scientific, San Jose, CA) for spike protein glycan analysis.

## Data and Materials availability

Mass Spectrometry data: ProteomeXchange Consortium via the PRIDE partner repository All data is available in the main text and supplemental information.

Information on recombinant proteins and reagents is available upon request

---

**Figure 2-figure supplement 1.**
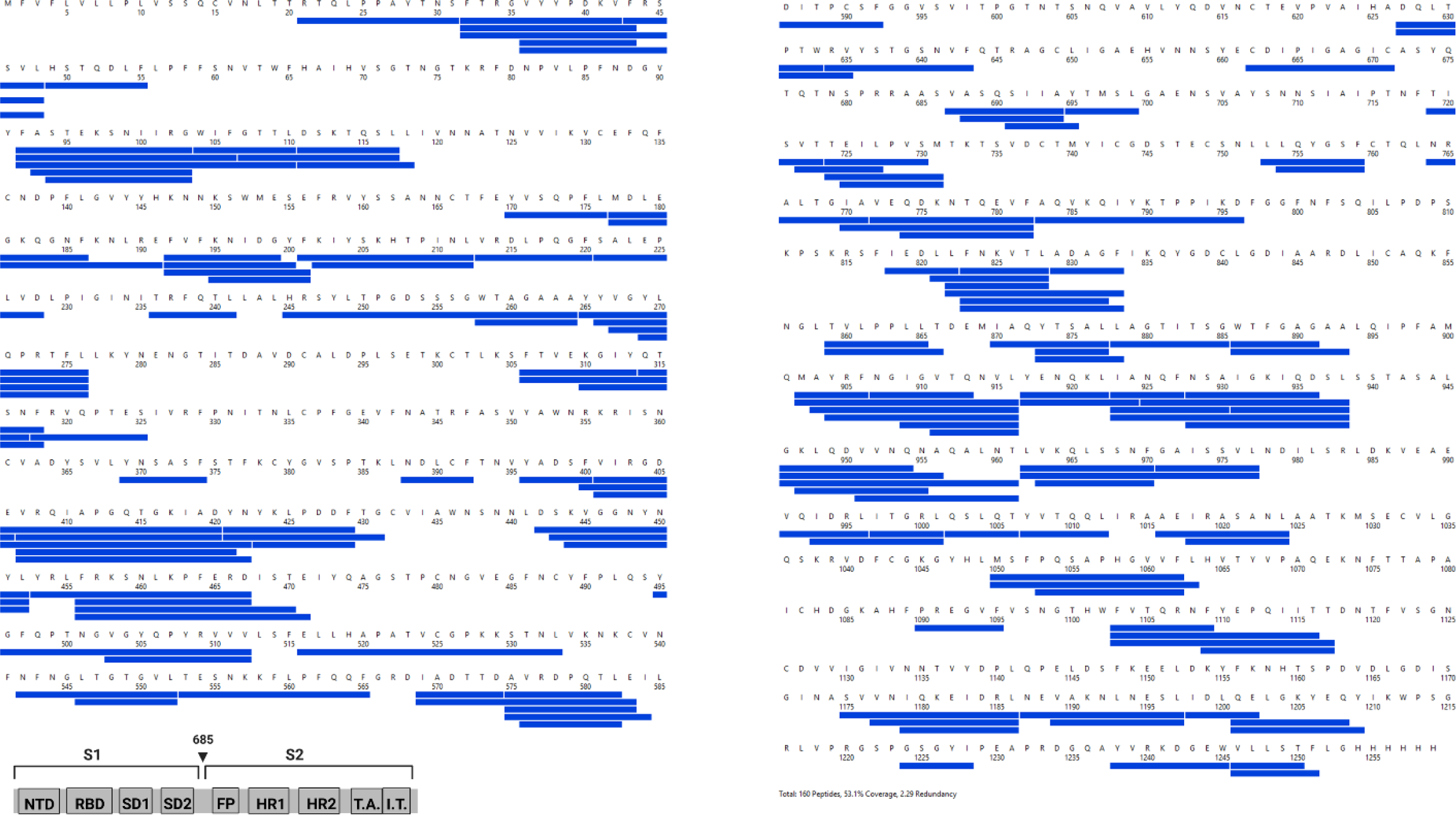
Primary sequence coverage for wildtype S 2P versus 6P comparison. Coverage map of wildtype 2P S compared to 6P S using the wildtype 2P sequence showing 160 peptides spanning 53.1% of the S protein. The domain organization of S is also shown.

**Figure 2-figure supplement 2.**
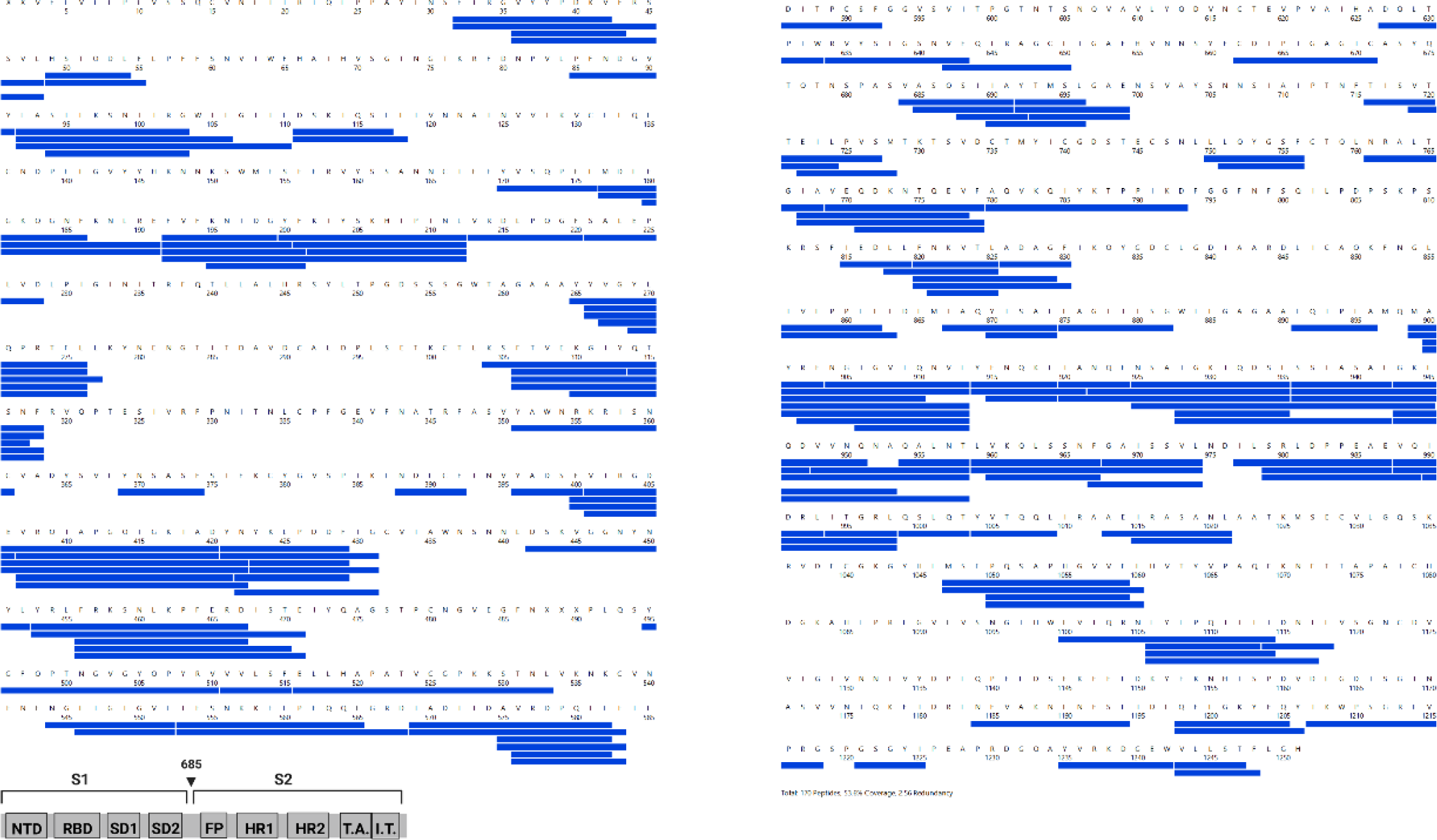
Primary sequence coverage for wildtype S incubated at 37°C versus unincubated wildtype S comparison. Coverage map of wildtype S incubated at 37°C compared to unincubated wildtype S using the wildtype 2P sequence showing 170 peptides spanning 53.3% of the S protein. The domain organization of S is also shown.

**Figure 2-figure supplement 3.**
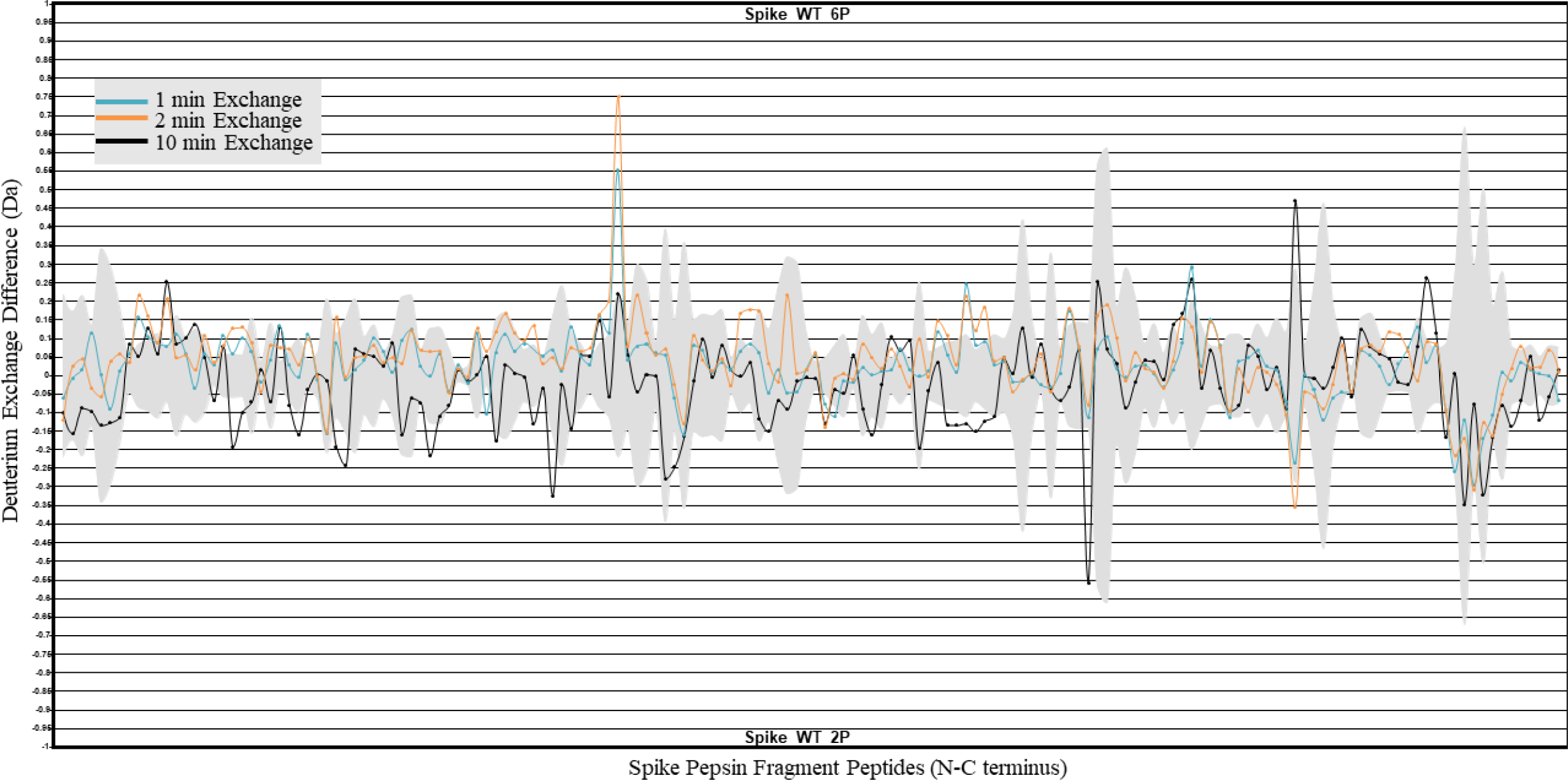
HDXMS analysis of 2P and 6P S constructs. Difference plot of Wildtype 6P S minus Wildtype 2P S for peptides N to C terminus. Differences for 1, 2, and 10 min exchange are shown in blue, orange, and black respectively. The grey trace denotes standard errors of deuterium exchange for each peptide.

**Figure 2-figure supplement 4.**
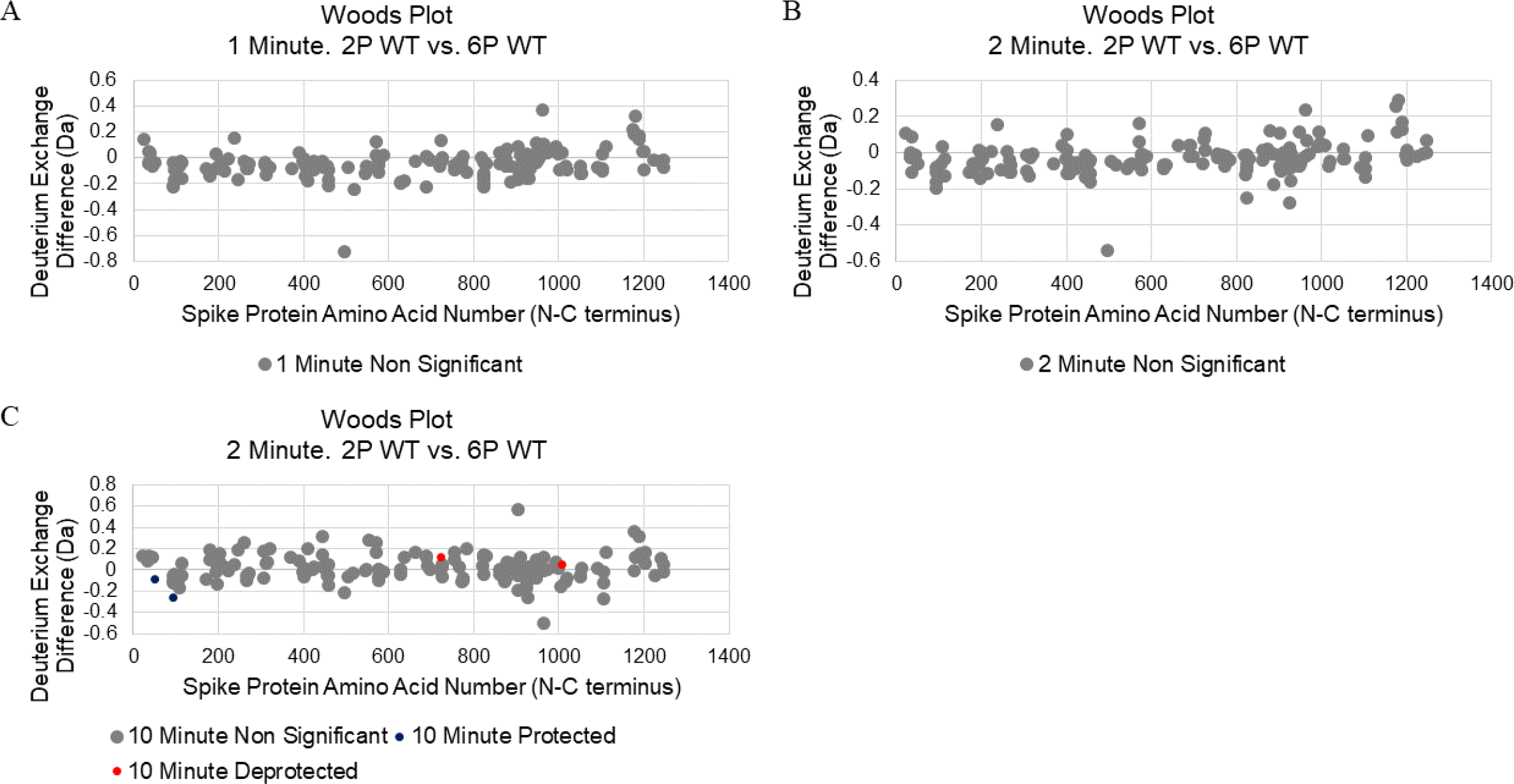
Woods plot analysis of Wildtype 2P versus 6P S. Woods plots for 1 min (A), 2 min (B), and 10 min (C) exchange. Significantly protected (blue) or deprotected (red) peptides were identified using a peptide level significance test and a P value <0.01.

**Figure 2-figure supplement 5.**
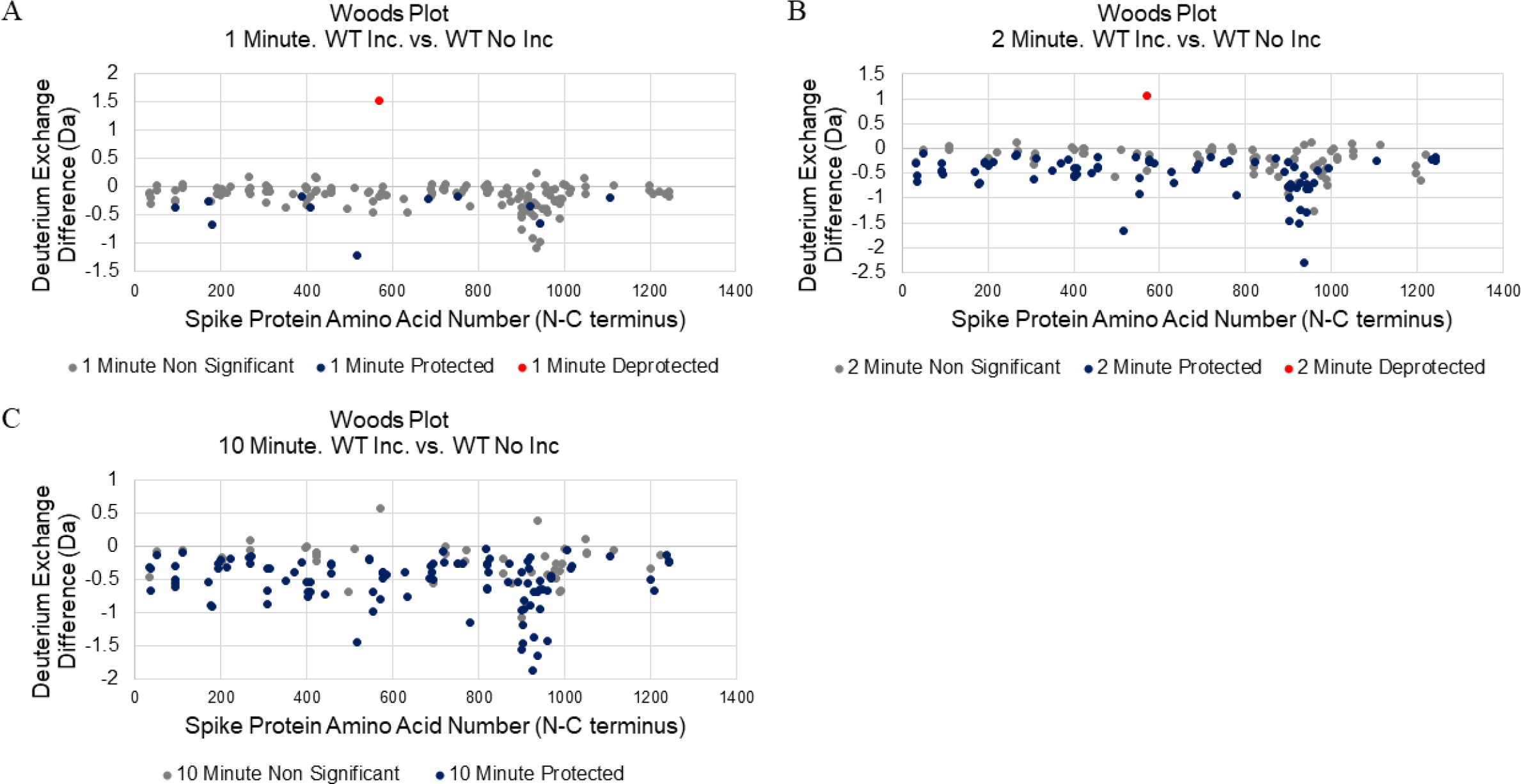
Woods plot analysis of wildtype S incubated at 37°C versus unincubated wildtype S. Woods plots for 1 min (A), 2 min (B), and 10 min (C) exchange. Significantly protected (blue) or deprotected (red) peptides were identified using a peptide level significance test and a P value <0.01.

**Figure 3-figure supplement 1.**
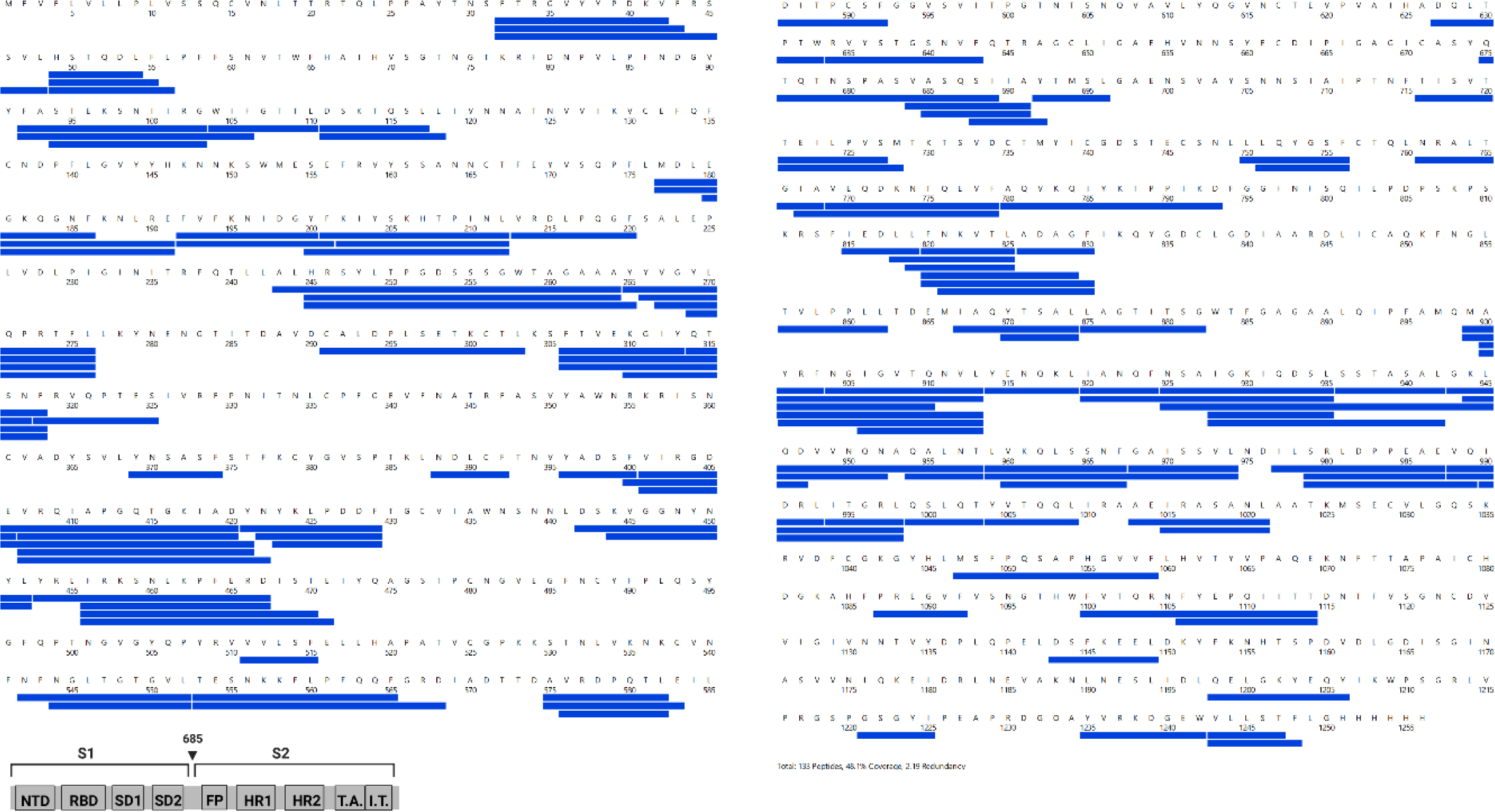
Primary sequence coverage for D614G S versus wildtype S comparison. Coverage map of D614G S compared to wildtype S using the Wildtype 2P sequence showing 132 peptides spanning 47.3% of the S protein. The domain organization of S is also shown.

**Figure 3-figure supplement 2.**
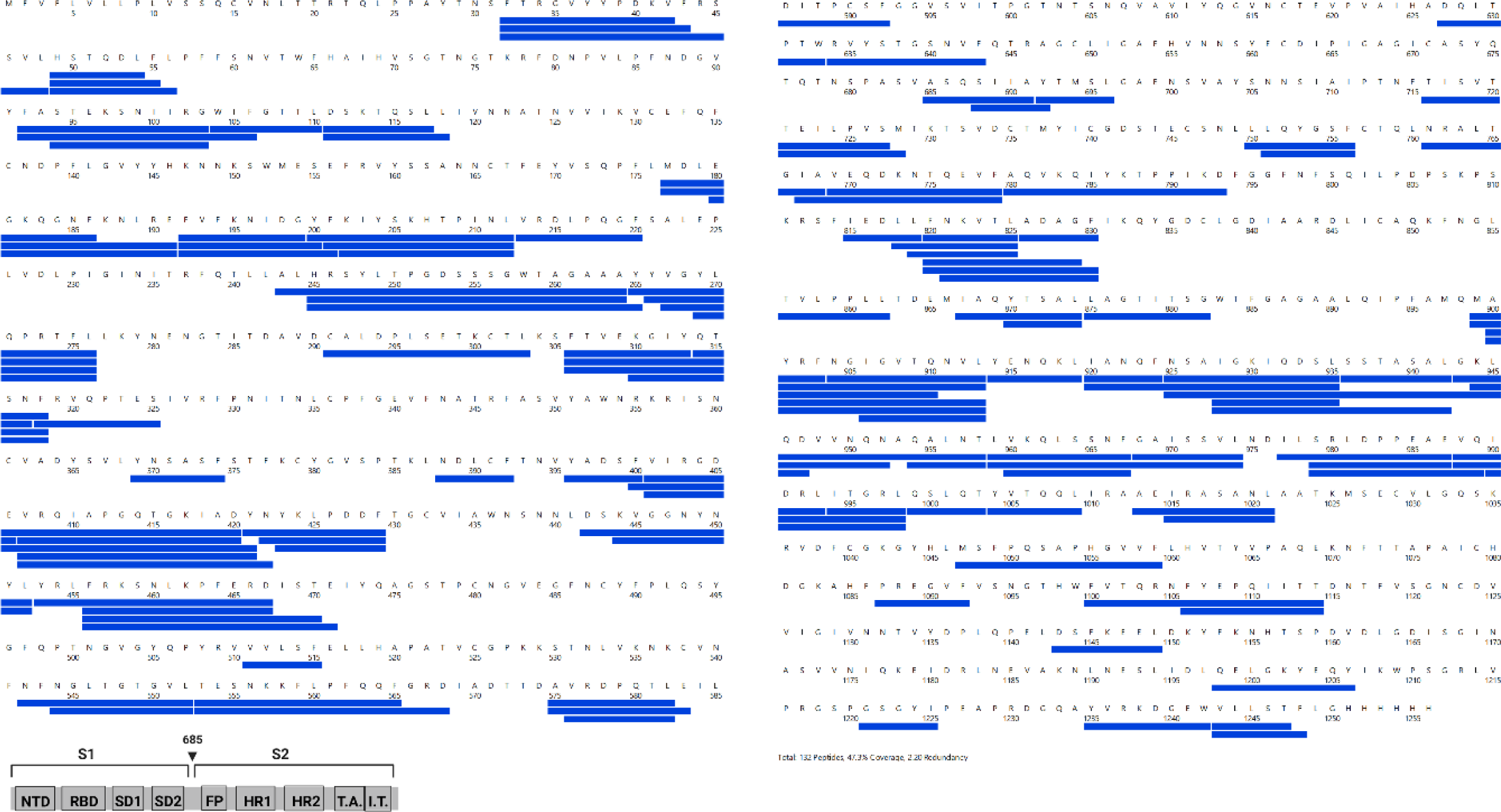
Primary sequence coverage for D614G S incubated at 37°C versus unincubated D614G S comparison. Coverage map of D614G S incubated at 37°C compared to unincubated D614G S using the D614G 2P sequence showing 133 peptides spanning 48.1% of the S protein. The domain organization of S is also shown.

**Figure 3-figure supplement 3.**
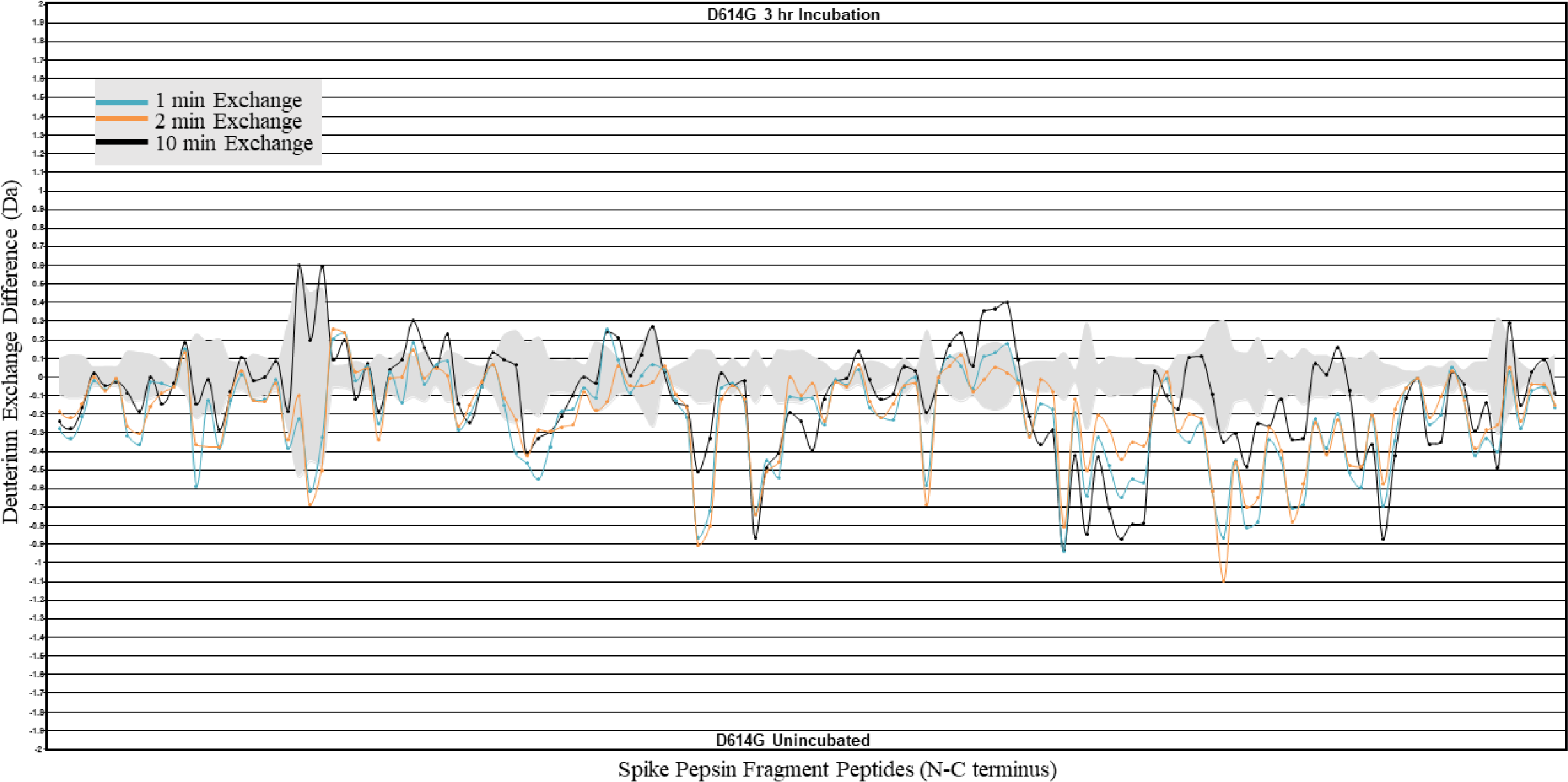
HDXMS analysis of incubation effects on D614G S. Difference plot of D614G S incubated for 3 h at 37 °C minus D614G S with no incubation for peptides N to C terminus. Differences for 1, 2, and 10 min exchange are shown in blue, orange, and black respectively. The grey trace denotes standard errors of deuterium exchange for each peptide.

**Figure 3-figure supplement 4.**
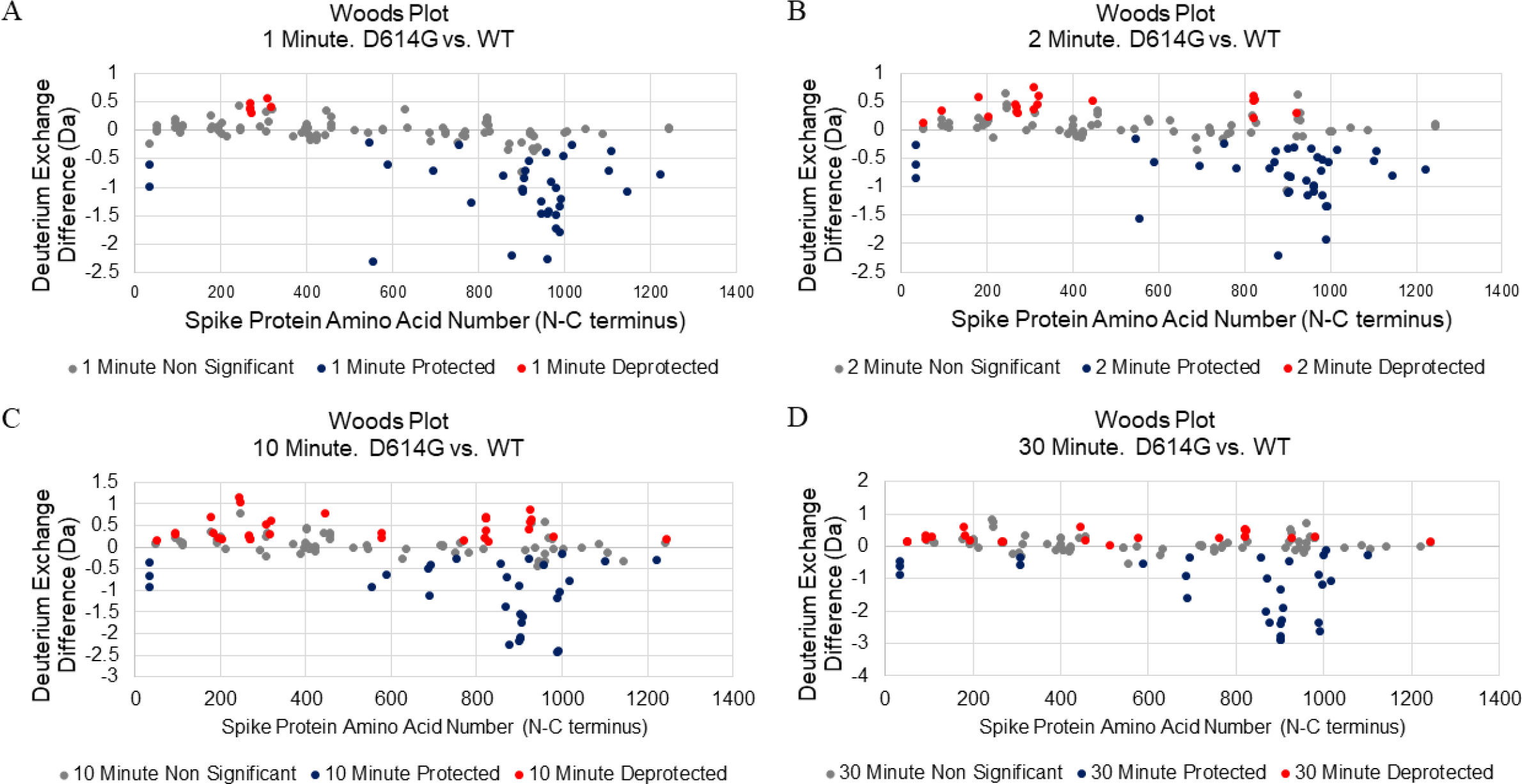
Woods plot analysis of D614G S versus Wildtype S. Woods plots for 1 min (A), 2 min (B), 10 min (C), and 30 min (D) exchange. Significantly protected (blue) or deprotected (red) peptides were identified using a peptide level significance test and a P value <0.01.

**Figure 3-figure supplement 5.**
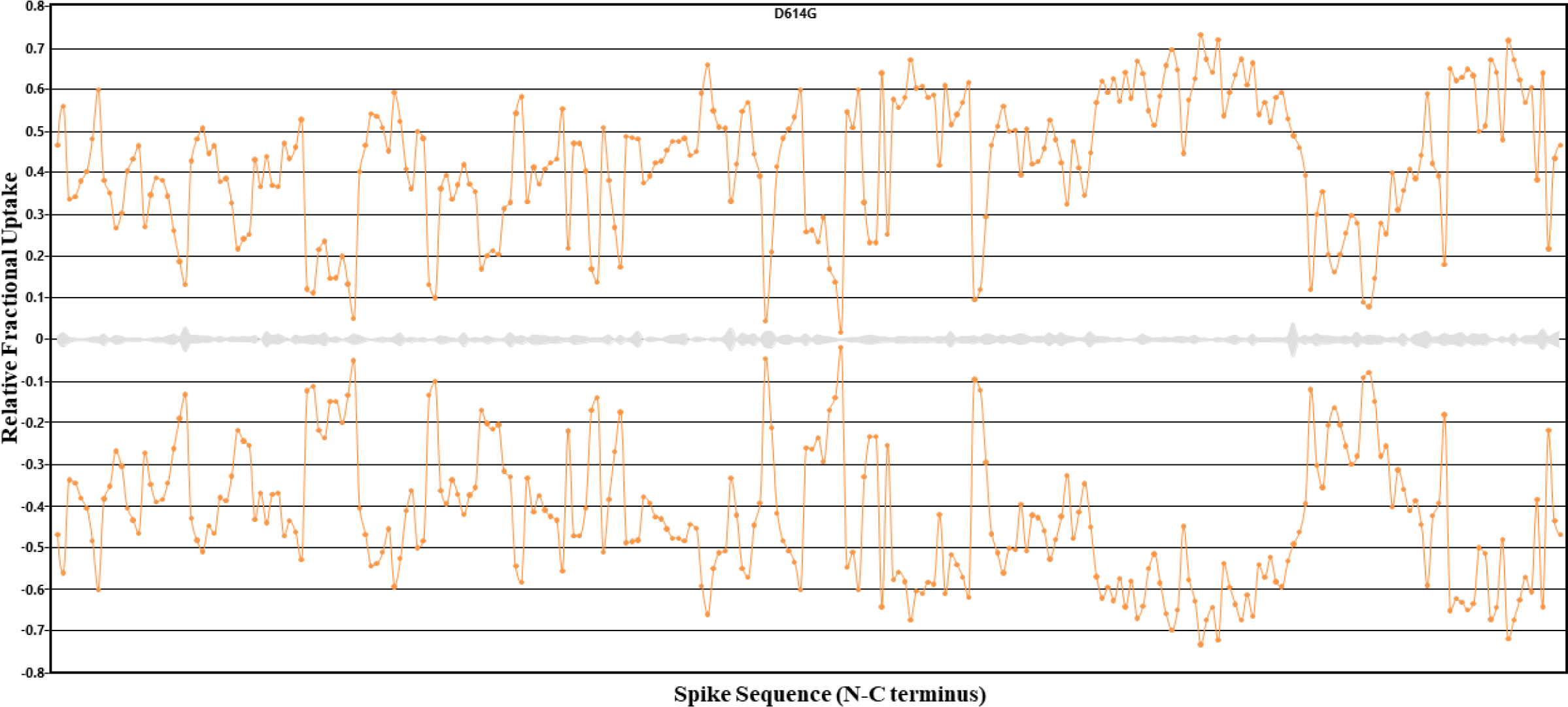
Back exchange measurements for pepsin fragment peptides from D614G S. Deuterium exchange in peptides after long deuteration (Dex = 24 h) is displayed as an RFU plot. The highest exchanging peptides (943-958 and 957-967) were used to calculate an average back exchange estimate of 20%.

**Figure 4-figure supplement 1.**
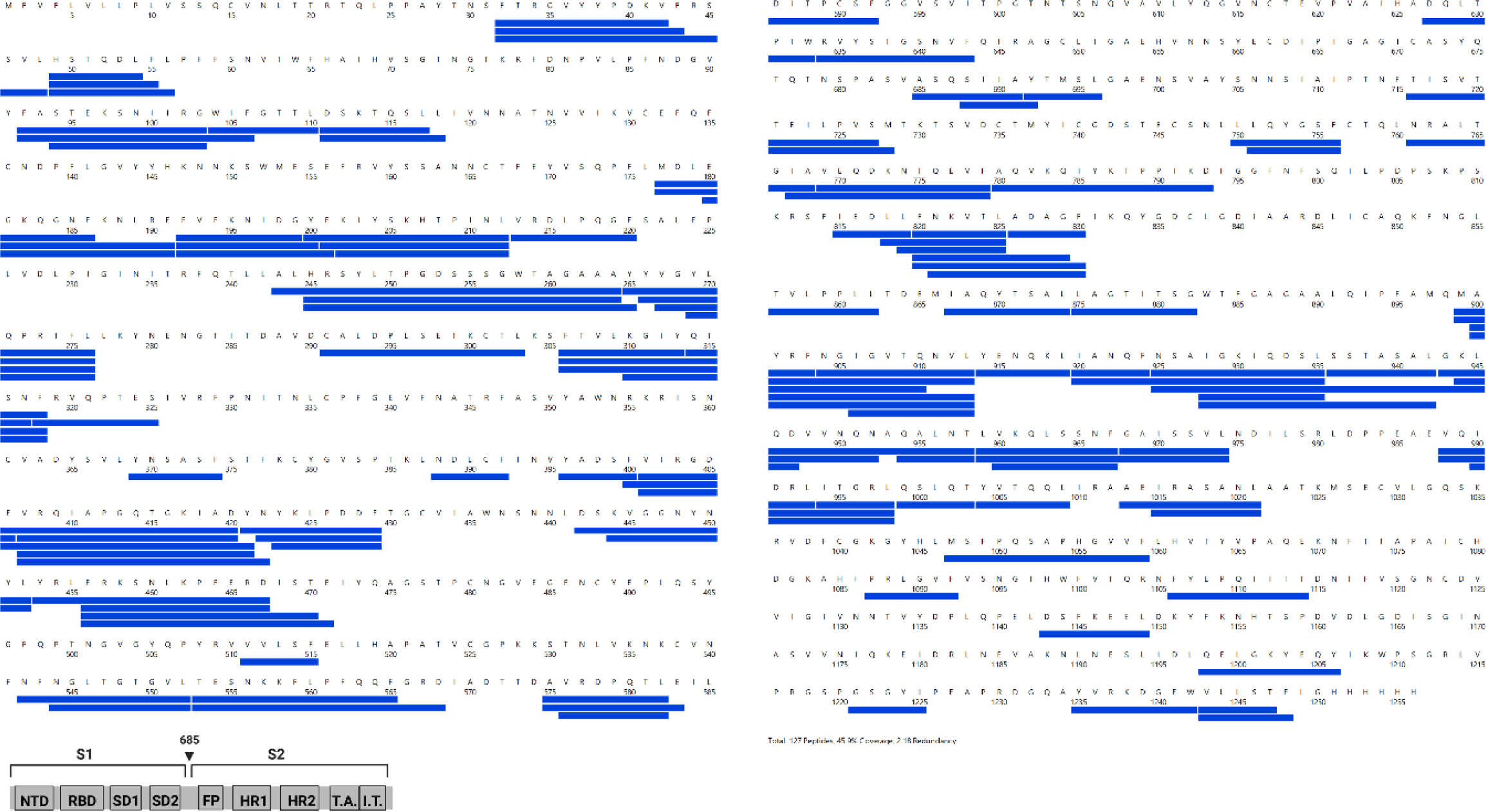
Primary sequence coverage for Alpha S versus D614G S comparison. Coverage map of Alpha S compared to D614G S using the D614G 2P sequence showing 127 peptides spanning 45.9% of the S protein. The domain organization of S is also shown.

**Figure 4-figure supplement 2.**
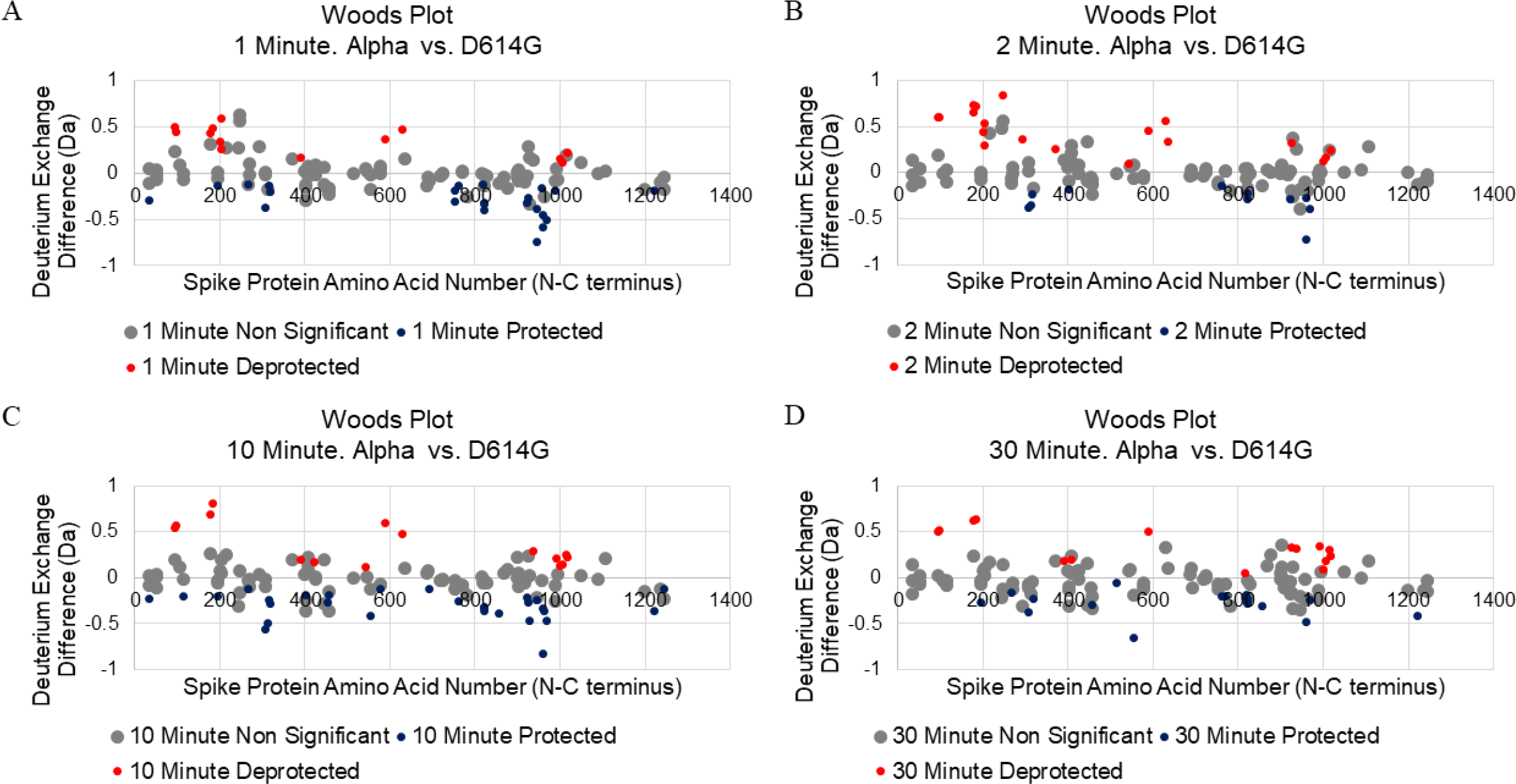
Woods plot analysis of Alpha variant S versus D614G S. Woods plots for 1 min (A), 2 min (B), 10 min (C), and 30 min (D) exchange. Significantly protected (blue) or deprotected (red) peptides were identified using a peptide level significance test and a P value <0.01.

**Figure 5-figure supplement 1.**
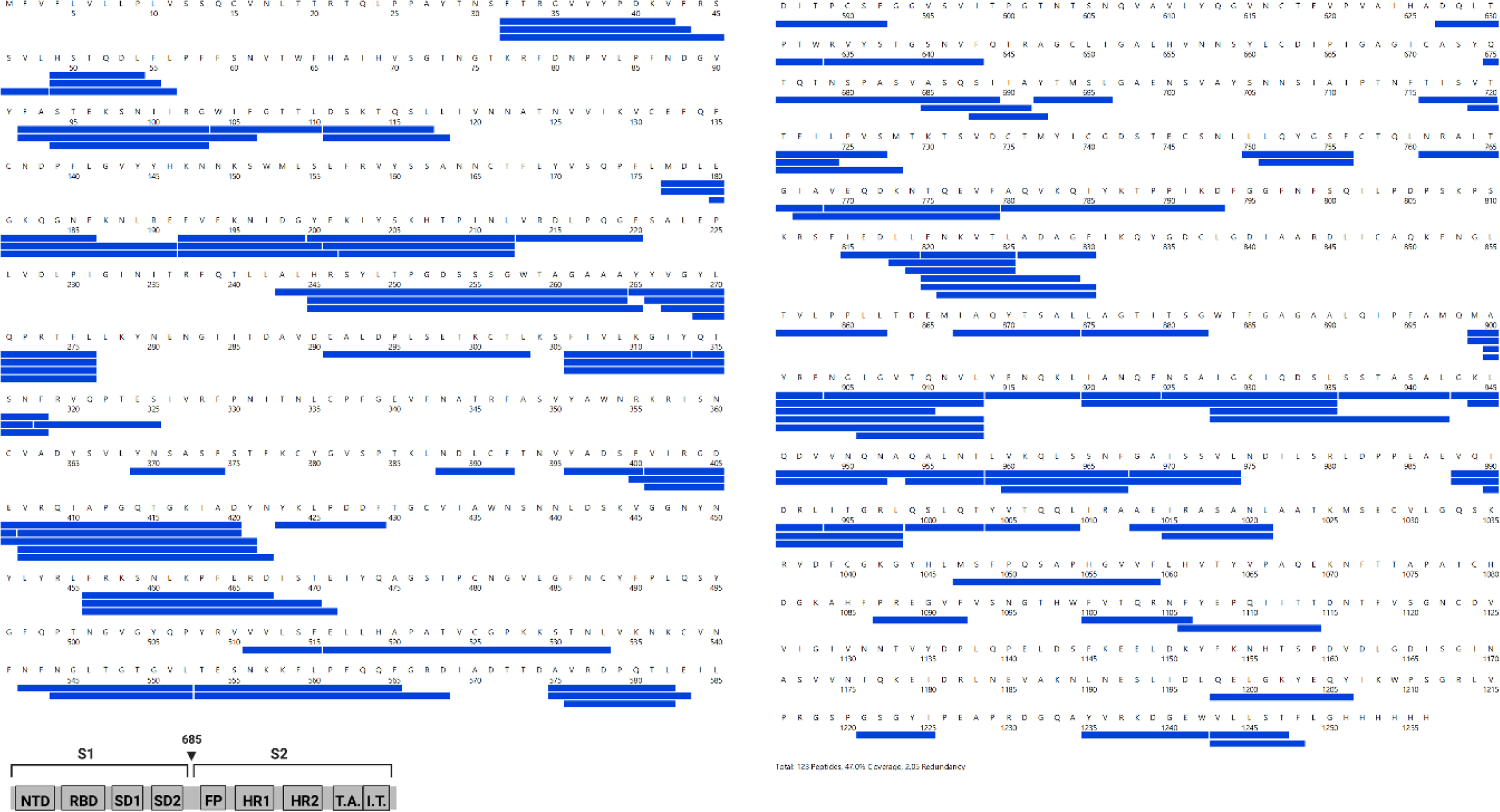
Primary sequence coverage for Delta S versus Alpha S comparison. Coverage map of Delta S compared to Alpha S using the D614G 2P sequence showing 123 peptides spanning 47.0% of the S protein. The domain organization of S is also shown.

**Figure 5-figure supplement 2.**
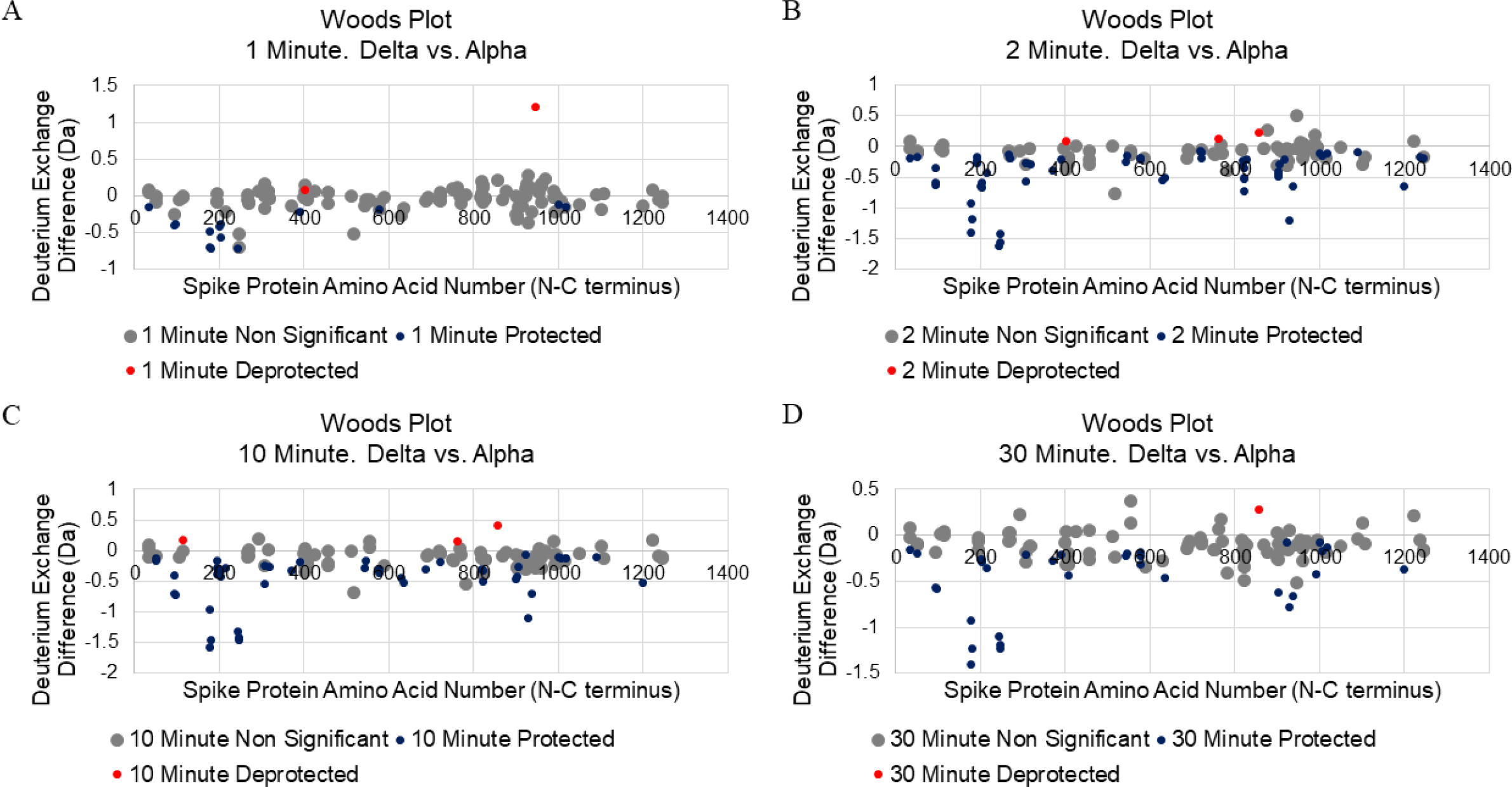
Woods plot analysis of Delta variant S versus Alpha variant S. Woods plots for 1 min (A), 2 min (B), 10 min (C), and 30 min (D) exchange. Significantly protected (blue) or deprotected (red) peptides were identified using a peptide level significance test and a P value <0.01.

**Figure 6-figure supplement 1.**
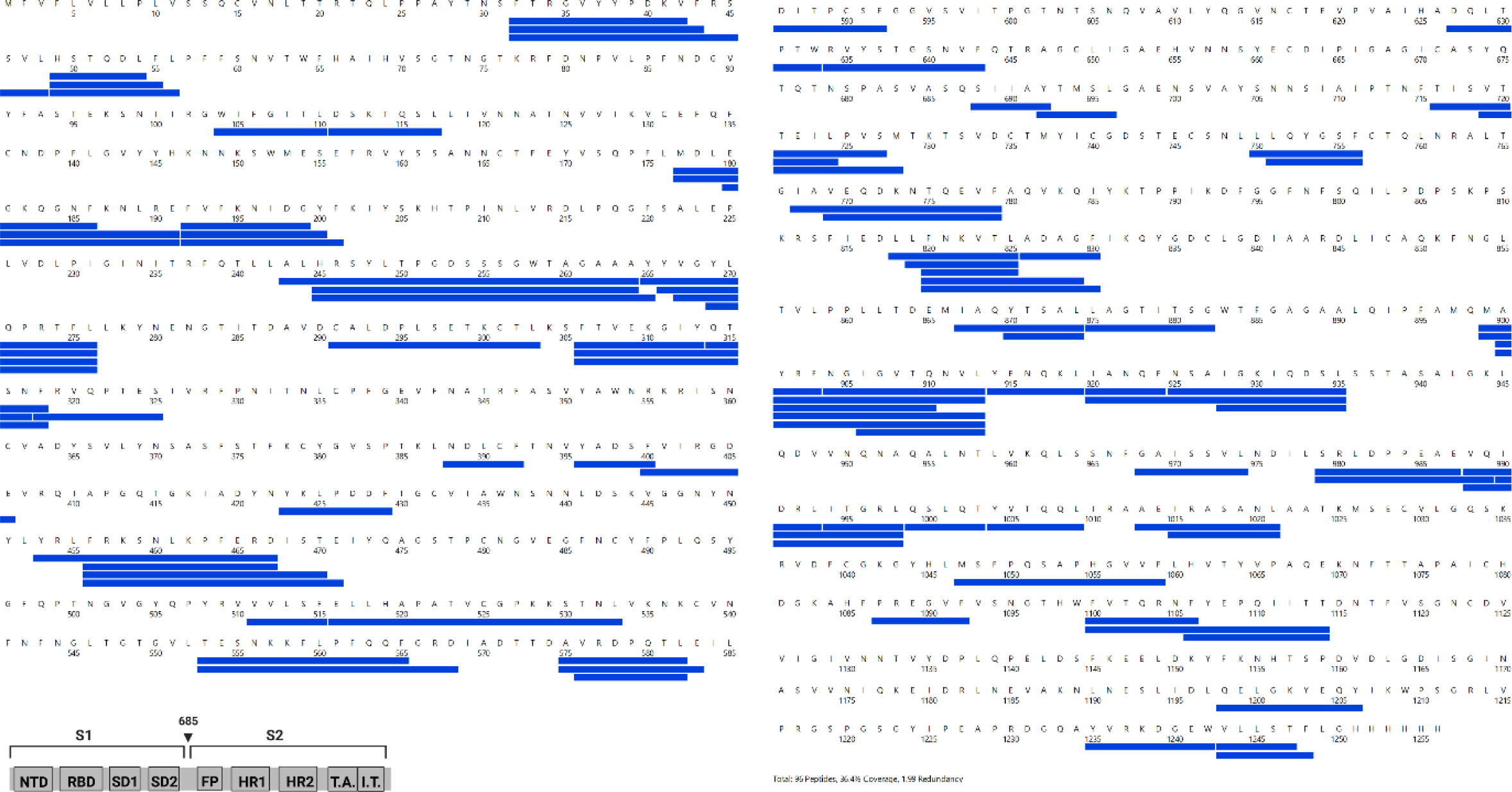
Primary sequence coverage for Omicron S versus Delta S comparison. Coverage map of Omicron S compared to Delta S using the D614G 2P sequence showing 96 peptides spanning 36.4% of the S protein. The domain organization of S is also shown.

**Figure 6-figure supplement 2.**
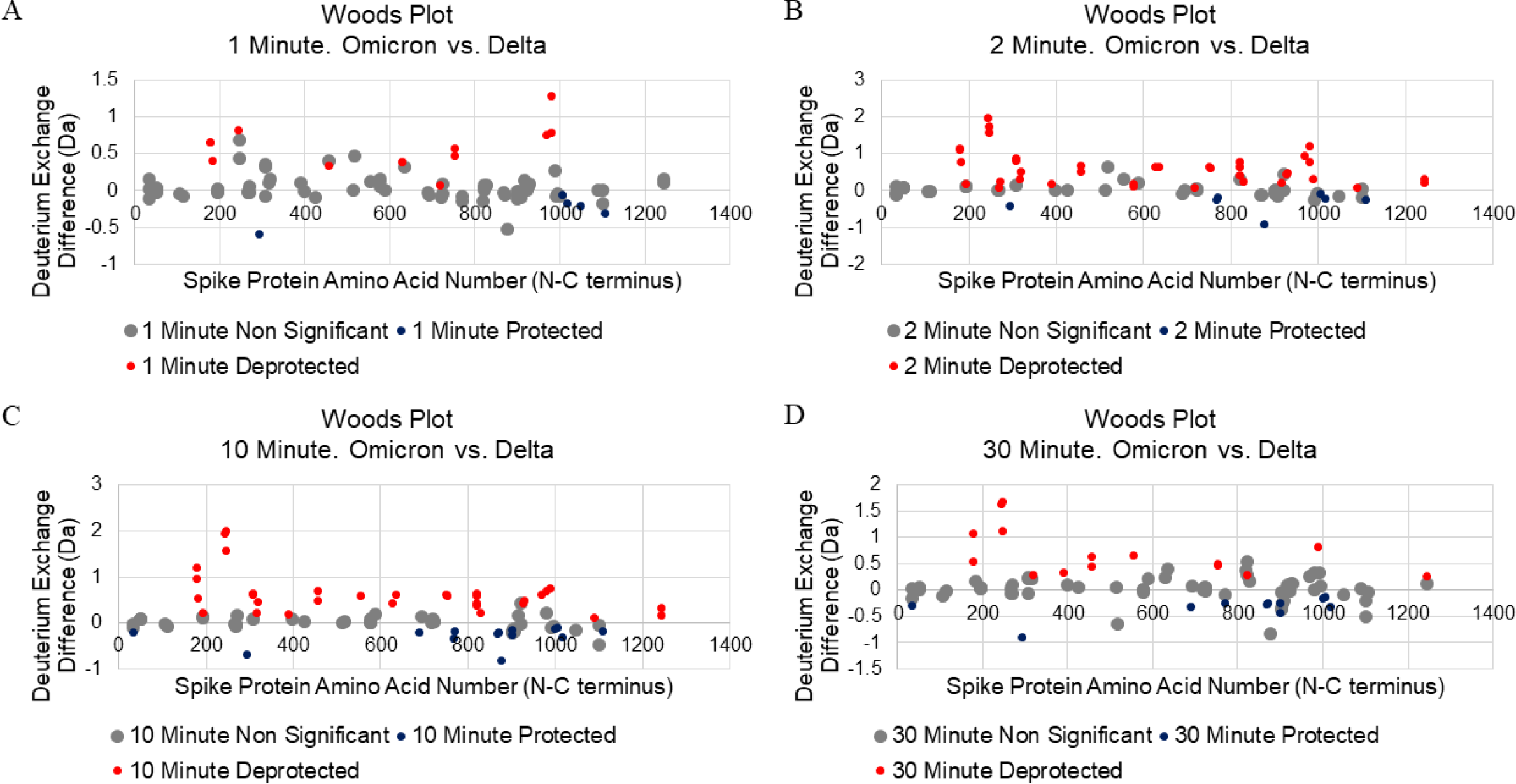
Woods plot analysis of Omicron variant S versus Delta variant S. Woods plots for 1 min (A), 2 min (B), 10 min (C), and 30 min (D) exchange. Significantly protected (blue) or deprotected (red) peptides were identified using a peptide level significance test and a P value <0.01.

**Figure 7-figure supplement 1.**
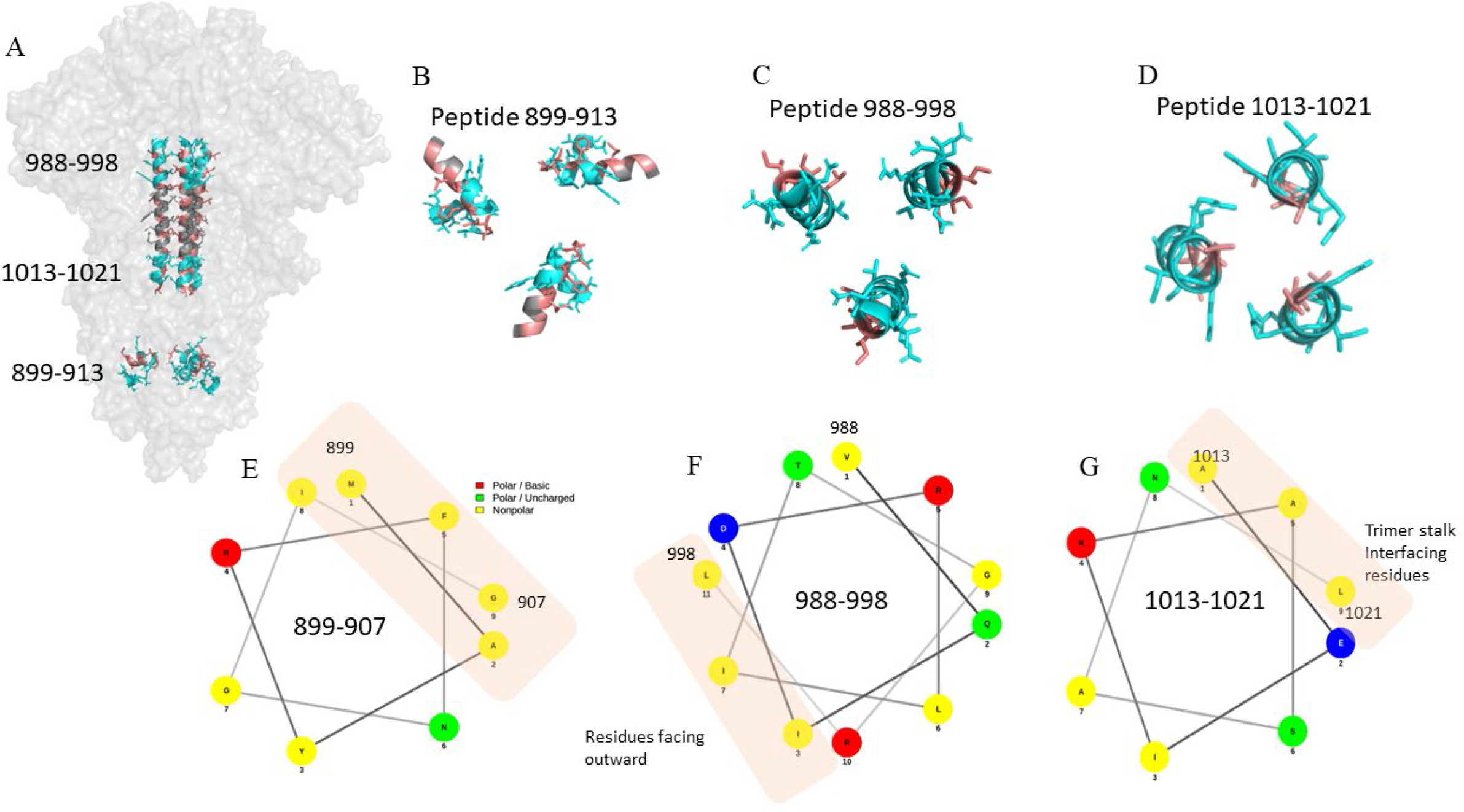
Hydrophobic interactions maintain trimer core at the stalk region. (A) Structure of the S trimer highlighting stalk region in salmon (PDB ID: 6VXX). HDXMS analysis peptides at the top, middle and bottom of the trimer stalk are colored cyan. (B, C, D) Cross-sectional view of peptides in the S trimer stalk are colored cyan. Hydrophobic residues are colored salmon. (E, F, G) Helical wheel representation of peptides classifying residues based on polarity. Hydrophobic patches are shown by orange rectangle.

**Figure 7-figure supplement 2.**
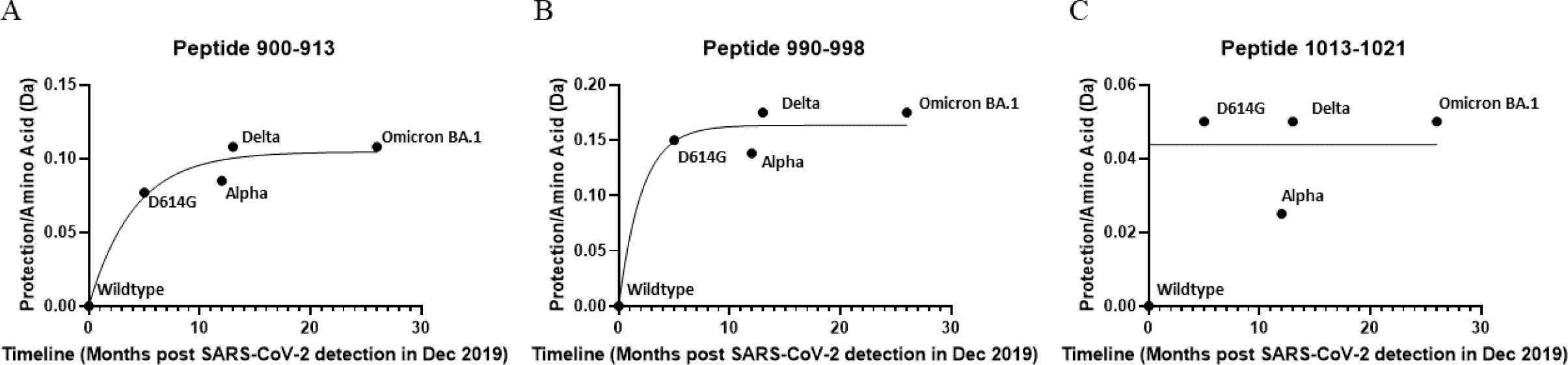
Plot of protection in S trimer stalk peptides as function of timeline of emergence. Protection for each peptide was determined by subtracting deuterium uptake for each variant from wildtype and normalizing to the number of exchangeable amino acids. Protection is plotted against months after the first identification of SARS-CoV-2 in Dec. 2019 and curves fit to a one phase association (Graphpad Prism 3.0, San Diego CA).

**Table S1.**
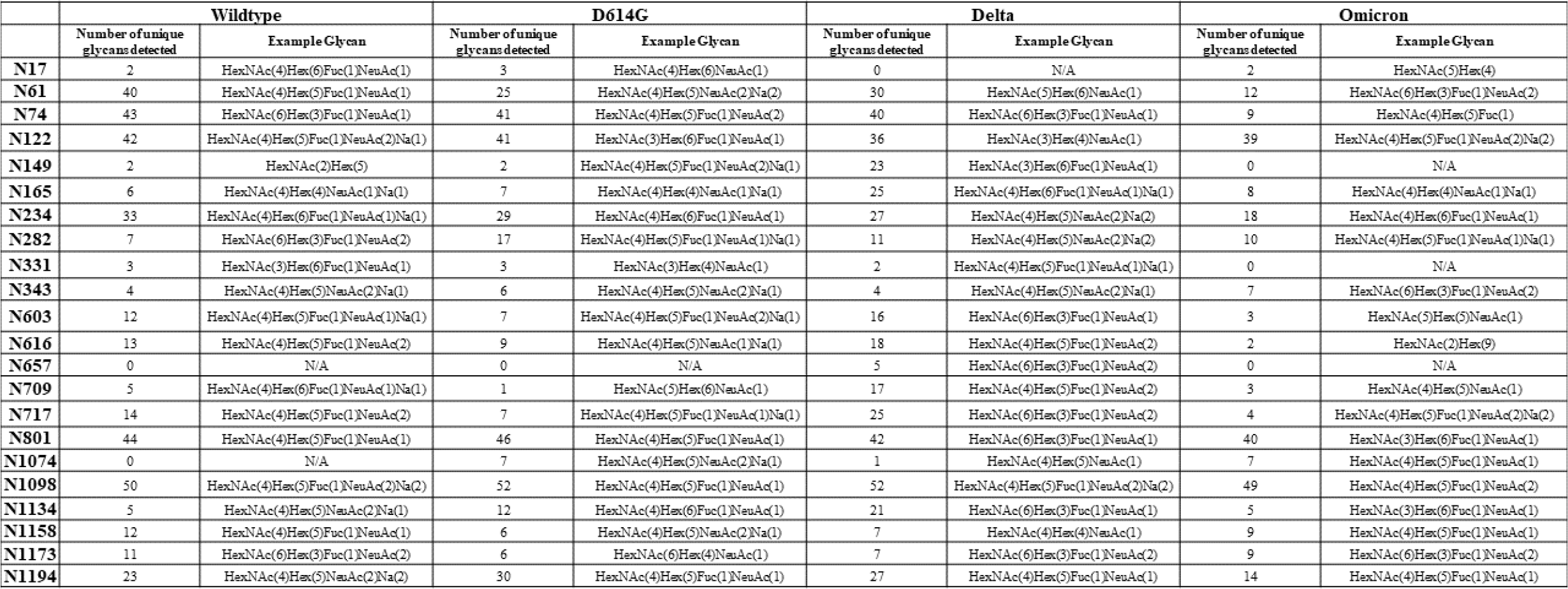
Glycosylation profile of SARS-CoV-2 spike variants. N-linked glycans were identified by mass spectrometry. The number of glycans identified at each site and an example glycan are reported.

**Table S2.** Differential ensemble behavior across S variants inferred from spectral broadening. A) Table showing spectral broadening at Dex = 1 min B) Table showing spectral broadening at Dex = 10 min. The number of spectral peaks were determined based on the assigned sticks during HDXMS analysis across common peptides.

| Number of spectral peaks in bimodal peptides at deuterium exchange time = 1 min |  |  |  |  |  |
| --- | --- | --- | --- | --- | --- |
| Peptide | WT | D614G | Alpha | Delta | Omicron (BA.1) |
| 553-568 | 15 | 13 | 13 | 13 | 13 |
| 875-882 | 7.5 ± 0.5 | 6 | 6.3 ± 0.57 | 5.6 ± 0.57 | 3.3 ± 0.57 |
| 900-913 | 12 | 10 | 10 | 9.6 ± 0.57 | 10 |
| 943-958 | 14.5 ± 0.5 | 13.3 ± 1.15 | 13 | 13.6 ± 0.57 | - |
| 988-998 | 9 | 6.3 ± 0.57 | 5 | 6.6 ± 0.57 | 6.3 ± 0.57 |

**B**
| Number of spectral peaks in bimodal peptides at deuterium exchange time = 10 min |  |  |  |  |  |
| --- | --- | --- | --- | --- | --- |
| Peptide | WT | D614G | Alpha | Delta | Omicron (BA.1) |
| 553-568 | 14 | 13 | 13 | 13 | 14 |
| 875-882 | 8 | 6.3 ± 0.57 | 7 | 6.3 ± 0.57 | 3 |
| 900-913 | 15 | 13 | 13 | 11 | 12 |
| 943-958 | 16 | 15.6 ± 0.57 | 16 | 12 | - |
| 988-998 | 10.3 ± 0.57 | 7.6 ± 1.15 | 5.3 ± 0.57 | 8±1 | 8 |

**Table S3.** Differences in NTD deuterium uptake comparing D614G and variants with WT at Dex = 30 min.

| Peptide | Sequence | D614G –<br>WT (ΔDex) | Alpha –<br>WT (ΔDex) | Delta –<br>WT<br>(ΔDex) | Omicron –<br>WT (ΔDex) |
| --- | --- | --- | --- | --- | --- |
| 177-191 | MDLEGKQGNFKNLRE | $0.5 \pm 0.2$ Da | $1.1 \pm 0.2$<br>Da | $-0.2 \pm 0.2$<br>Da | $0.3 \pm 0.2$ Da |
| 245-265 | HRSYLTPGDSSSGWTAGAAAY | $0.7 \pm 0.4$ Da | $0.6 \pm 0.4$<br>Da | $-0.6 \pm 0.5$<br>Da | $1.1 \pm 0.3$ Da |
| 456-467 | FRKSNLKPFERD | $0.2 \pm 0.1$ Da | $0.0 \pm 0.1$<br>Da | $0.0 \pm 0.1$<br>Da | $0.5 \pm 0.1$ Da |

**Table S4.**
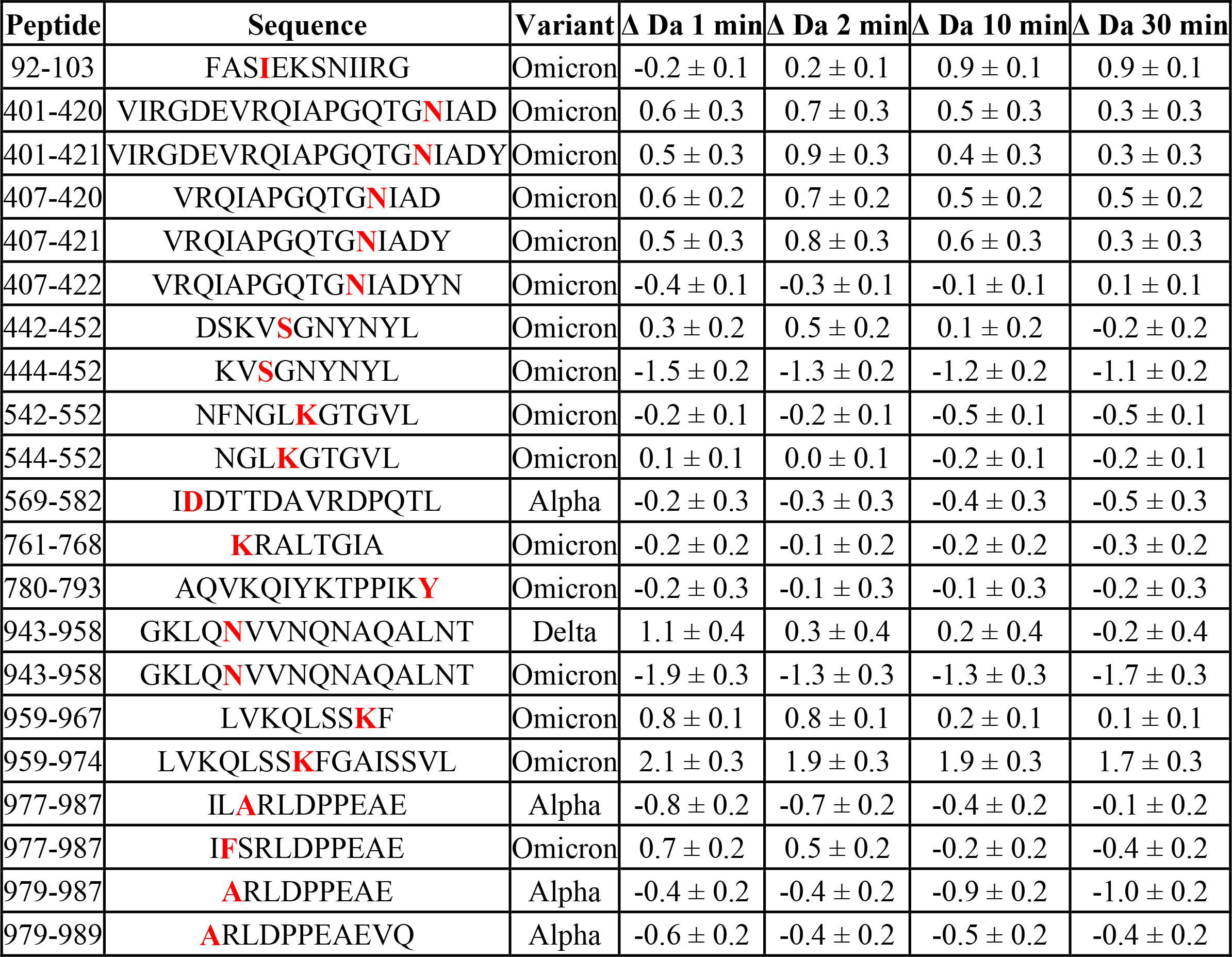
HDXMS analysis of mutated peptides. Differences between variants and D614G S for mutated peptides reported in Da for 1, 2, 10, and 30 min exchange. Mutations sites are shown in bold red. Type or paste caption here.

